# Network reconstruction for trans acting genetic loci using multi-omics data and prior information

**DOI:** 10.1101/2020.05.19.101592

**Authors:** Johann S. Hawe, Ashis Saha, Melanie Waldenberger, Sonja Kunze, Simone Wahl, Martina Müller-Nurasyid, Holger Prokisch, Harald Grallert, Christian Herder, Annette Peters, Konstantin Strauch, Fabian J. Theis, Christian Gieger, John Chambers, Alexis Battle, Matthias Heinig

## Abstract

**Background:** Molecular multi-omics data provide an in-depth view on biological systems, and their integration is crucial to gain insights in complex regulatory processes. These data can be used to explain disease related genetic variants by linking them to intermediate molecular traits (quantitative trait loci, QTL). Molecular networks regulating cellular processes leave footprints in QTL results as so-called *trans* -QTL hotspots. Reconstructing these networks is a complex endeavor and use of biological prior information has been proposed to alleviate network inference. However, previous efforts were limited in the types of priors used or have only been applied to model systems. In this study, we reconstruct the regulatory networks underlying *trans* -QTL hotspots using human cohort data and data-driven prior information.

**Results:** We devised a strategy to integrate QTL with human population scale multi-omics data and comprehensively curated prior information from large-scale biological databases. State-of-the art network inference methods applied to these data and priors were used to recover the regulatory networks underlying *trans* -QTL hotspots. We benchmarked inference methods and showed, that Bayesian strategies using biologically-informed priors outperform methods without prior data in simulated data and show better replication across datasets. Application of our approach to human cohort data highlighted two novel regulatory networks related to schizophrenia and lean body mass for which we generated novel functional hypotheses.

**Conclusion:** We demonstrate, that existing biological knowledge can be leveraged for the integrative analysis of networks underlying *trans* associations to deduce novel hypotheses on cell regulatory mechanisms.

## Background

Genome-wide associations studies (GWAS) have been tremendously successful in discovering disease associated genetic loci. However, establishing causality or obtaining functional explanations for GWAS SNPs is still challenging. In recent years, the focus has shifted from discovery of disease loci to mechanism and explanation, and large efforts have been put into unravelling the functional consequences of GWAS SNPs [1, 2]. These have been made possible through technological advances in measuring genome-wide molecular data in large population cohorts, which further led to a steady increase in biological resources providing simultaneous measurements of different molecular layers (often termed *multi-omics* data). To elucidate disease mechanisms, systems genetics approaches seek to link GWAS SNPs to intermediate molecular traits by identifying quantitative trait loci (QTL) [3, 4], for example for gene expression levels (eQTL) [5–7] or DNA methylation at CpG dinucleotides (meQTL) [8–10].

Genetic variants that are QTL for quantitative molecular phenotypes that reside on a different chromosome are called *trans* -QTL. Previously, *trans* -QTL studies were successful in model systems [11, 12]. Recently, large-scale meta analyses of molecular QTL in very large sample sizes have now been applied to successfully map large numbers of *trans* -QTL in humans [7]. These are particularly interesting, as they have been found to be enriched for disease associations [7, 8, 13]. Yet, the underlying mechanisms leading to such associations can usually not be explained in a straightforward way [6], and in fact, 83% of discovered *trans* -eQTL in human are estimated to still be unexplained [7].

*Trans*-QTL hotspots [14], where a single genetic locus influences numerous quantitative traits on different chromosomes, can be seen as footprints of regulatory molecular networks and likely encode master regulators. One way of mechanistically explaining the effects of these master regulators is by reverse engineering the regulatory networks, and hence determining the intermediate molecular processes giving rise to the observed *trans* effects, ultimately yielding novel insights into disease pathophysiology [1, 14–16].

A large body of work has focused on inferring regulatory interactions from high-throughput data by individually combining distinct genomic layers like gene expression levels and genotype [6, 17–19] or chromosomal aberration [20] information. Generally, network inference to uncover regulatory mechanisms in biological systems has gotten much interest [15, 21–24]. The emergence of multi-omics data now also allows for establishing networks across more than two omics layers in a holistic approach to obtain more insight into the function of regulatory elements [16]. Major efforts have been made to recover functional interactions from such data, but methods to successfully reverse engineer regulatory networks across multiple omics layers are still lacking [1, 4, 25, 26].

Furthermore, utilizing the wealth of data available from genomic databases as biological prior information can guide the inference of complex multi-omics networks [26–28]. For instance, using known relationships discovered in previous studies as prior knowledge, such as protein-protein interactions (PPIs) or eQTL, can facilitate network reconstruction on novel datasets. Application of priors has been investigated in numerous works [e.g. 15, 27, 29–34], and while several studies show the advantage of using priors in synthetic datasets [22, 31, 33, 34] or model systems [15, 32, 34, 35], relatively few studies apply their inference methodologies to functional genomics data in humans [29, 33, 36, 37]. In case human data is considered, either cell line data are used [36], the inference is restricted to a single pathway [37] or no informative priors are used for this specific context [29]. Zuo *et al.* apply prior based inference to human cancer gene expression data, however, they only use priors based on PPIs extracted from the STRING database and focus on differential expression analysis [33]. What is still missing, is, to comprehensively integrate the vast amount of functional data from large-scale databases [38–41] as prior information in human multi-omic *trans* -QTL studies and to determine the appropriate inference methods.

Here, we developed a novel approach for understanding the molecular mechanisms underlying the statistical associations of *trans* -QTL hotspots by integrating existing biological knowledge and available multi-omics data to infer regulatory networks. We derived a comprehensive set of continuous priors from public datasets such as GTEx, the BioGrid and Roadmap Epigenomics and applied state-of-the-art network inference methods including graphical lasso [42], BDgraph [29] and iRafnet [32], and showed, that methods using data-driven priors outperform non-prior approaches for network reconstruction on simulated data. Moreover, we showed that networks inferred on real-world data using priors can be replicated more faithfully across independent datasets than networks inferred without priors. Finally, we demonstrated, that incorporating existing knowledge with multi-omics data yields novel insights into disease related cellular mechanisms when applied to real-world population cohort data of different omics types and tissues.

## Results

### Trans-QTL hotspots define regulatory network candidates

In this study, we aimed to reconstruct regulatory networks to explain *trans* quantitative trait locus (*trans* -QTL) hotspots on a molecular level through simultaneous integration of multiomics data [4]. *Trans*-QTL hotspots have previously been associated with disease [8, 13], and understanding their mechanisms of action can deepen our insights into regulatory pathways and, ultimately, into the disease process.

Our general analysis strategy is depicted in Figure 1A and consists of the following steps: 1) curate QTL hotspots, 2) gather functional data and prior information, 3a+b) benchmark network inference methods in simulation and replication study to select best suited method and 4) infer and interpret networks identified in the cohort data.

**Figure 1:**
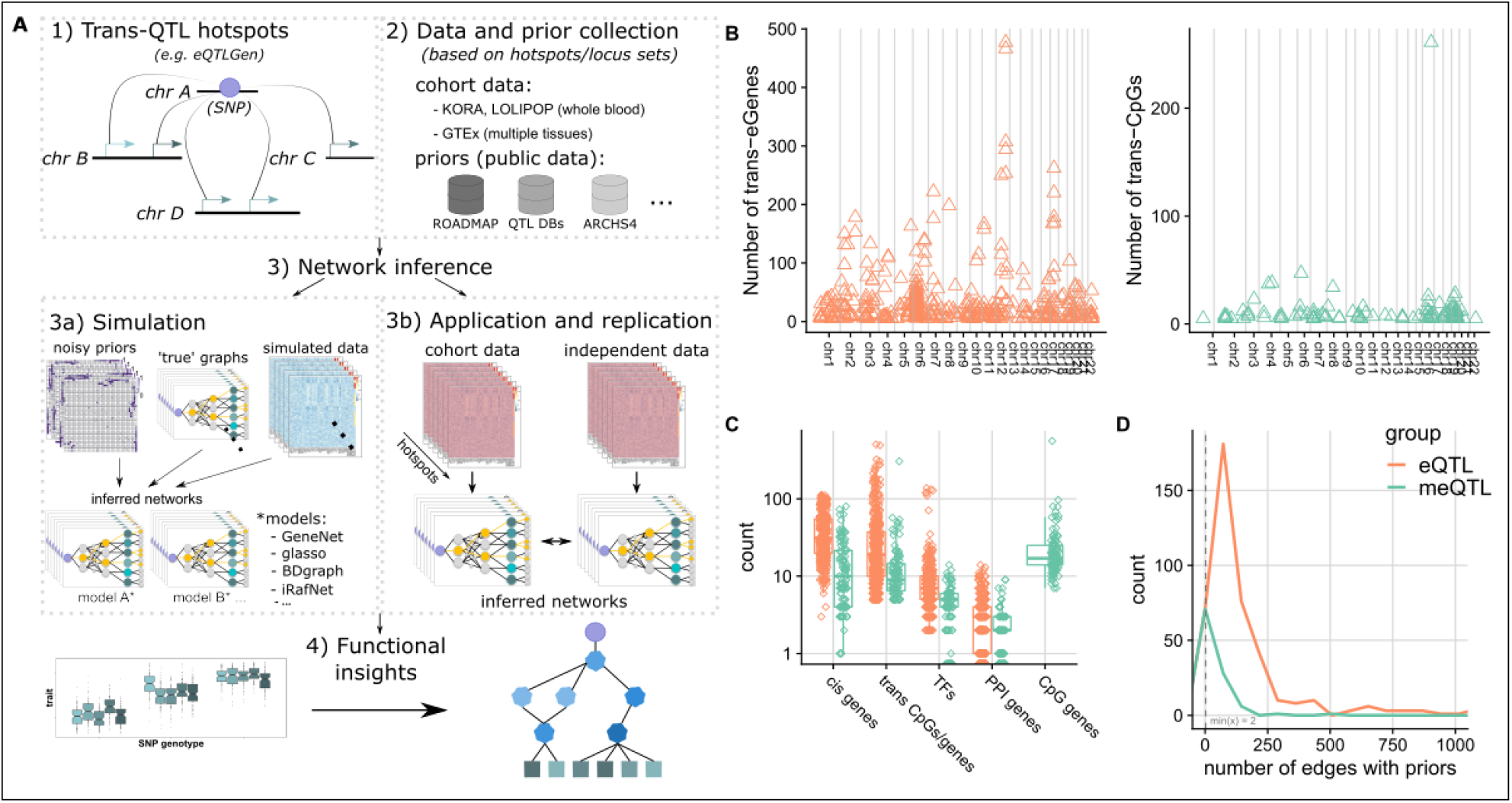
Project overview. Panel **A**) shows a graphical abstract of the analyses performed in this project. Panel **B**) provides a global view on the collected eQTL (orange) and meQTL (green) hotspots. The x-axis indicates ordered chromosomal positions for *trans* eGenes and CpG sites, respectively. Panel **C**) shows the total number of different genomic entities gathered over all hotspots during locus set creation (log scale). Panel **D**) depicts density plots of the number of possible network edges with available prior information (x-axis) over all hotspots, zoomed in to area between 0 and 1000. Same color coding is used in panels **B-D**.

We obtained *trans* hotspots from the methylation QTL (meQTL) discovered in whole-blood in the KORA [43] and LOLIPOP [44] cohorts reported by Hawe and colleagues [10] and the expression QTL (eQTL) published by the eQTLGen consortium [7], yielding a total of 107 and 444 *trans* -loci per QTL type, respectively (Figure 1B, see Methods for details). In addition to the whole-blood derived hotspots, we curated a single *trans* -eQTL hotspot in Skeletal Muscle tissue from GTEx v8 [38, 39], which we analyzed separately.

For each hotspot, we aimed to identify the causal gene at the genetic locus affected by the SNP and the intermediate genes which mediate the observed *trans* associations. To this end, we collected sets of candidate genes with different roles for each locus, which we term ‘locus sets’ (see Methods). A locus set contains the SNP defining the hotspot, the respective *trans* associated traits (CpGs for meQTL and genes for eQTL, ‘eGenes’), *cis* genes encoded near the SNP as candidate causal genes, *trans* genes (for meQTLs, genes in vicinity of the CpGs), as well as transcription factors (TFs) binding near the *trans* associated entities and PPI genes residing on the shortest path between *trans* traits and *cis* genes in a protein-protein interaction (PPI) network, as potential intermediate genes. *Cis* genes form potential candidate regulator genes of the locus, and the inclusion of the PPI and TF binding information allows us to bridge the inter-chromosomal gap between the SNP and the *trans* CpG sites/*trans* eGenes. An overview of entities collected over all loci for both QTL types is given in Figure 1C.

One main aspect of this work is the use of any form of biological prior information, including continuous scores, to guide network inference. We hence collect prior information for all possible edges between entities contained in locus sets in addition to the functional data (Figure 1). In total, four distinct types of edges are annotated with prior information: *SNP-Gene, Gene-Gene, TF-CpG/TF-Gene* and *CpG-Gene* edges. All prior information is generated from matched, public data independent of the data used during network inference (see Methods for details).

Figure 1D indicates the total number of edges annotated with prior information over all hotspots. For meQTL and eQTL, a minimum of 2 and 3 edges per hotspot show prior evidence, respectively, and most hotspots get only relatively few priors compared to the total number of possible edges (median 26 and 94, respectively). However, in both cases several networks collect priors for over 100 edges (8 and 209 loci with >= 100 priors for meQTL and eQTL). As expected, the total number of edges with prior information per locus correlates with the total number of possible edges in the respective loci, however, the fraction of all possible edges annotated with prior information decreases (Additional File 1, Figure S2).

### Benchmark of network inference methods

#### Simulation study shows benefit of data-driven priors

Numerous methods for regulatory network inference have been proposed (e.g. [42, 45, 46], see also [4]), and, therefore, before investigating individual hotspots in detail we sought to select the method best suited for this study (see Figure 1A step 3). To this end, we performed an extensive simulation study (Figure 1A step 3a) to evaluate the performance of five distinct methodologies (see Table 1 for a method overview) in reconstructing ground truth graphs from simulated data and prior information. Simulated data were matched with the observed QTL-hotspots by preserving the sample size and the total number of input nodes and 100 simulations were performed for each hotspot. We evaluated the impact of priors for different sample sizes by sub-sampling the simulated data and using the full prior matrix. To assess the impact of noise in priors, we inferred networks separately from prior information with varying degrees of noise (up to 100%, see Methods for details) for the complete data.

**Table 1:**
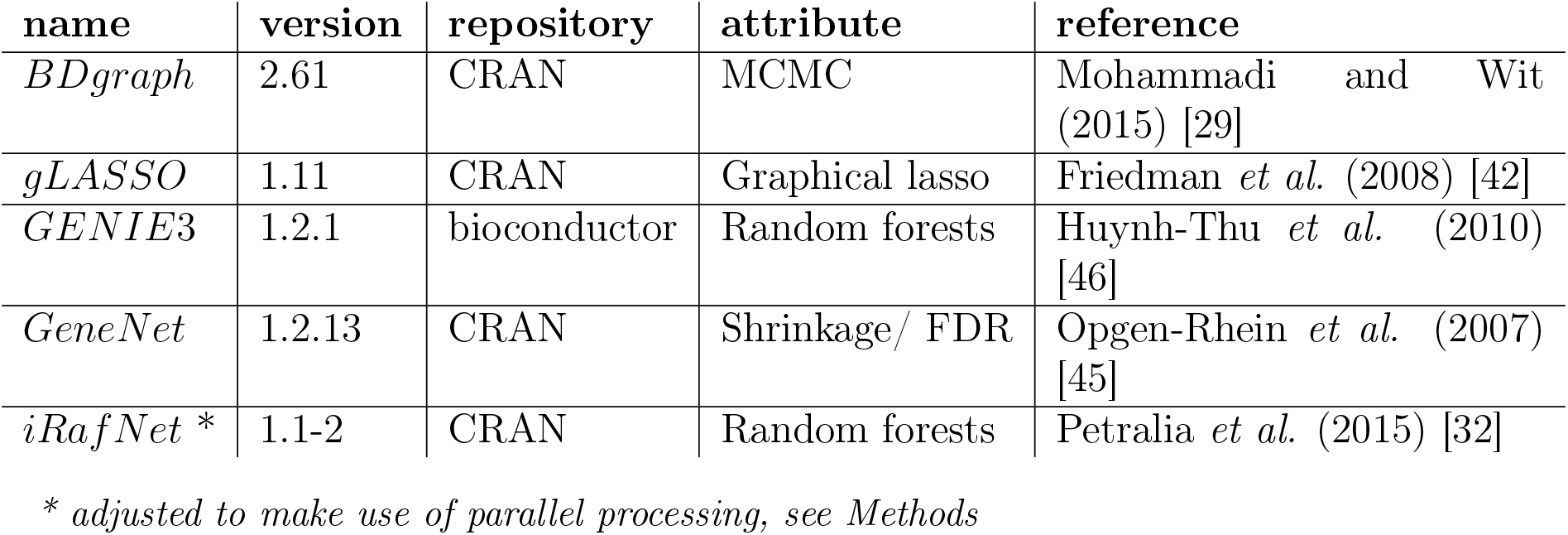
Overview of the network inference packages used in the simulation study. * *adjusted to make use of parallel processing, see Methods*

We gauge the relative gain in performance attributable to prior information for both *gLASSO* and *BDgraph* by always training two distinct models, one utilizing the provided priors (*gLASSO_P_*, *BDgraph_P_*) and one without priors *(gLASSO*, *BDgraph*). The implementation of *iRafNet* always requires a prior matrix, whereas both *GeneNet* and *GENIE*3 cannot utilize prior information and hence were trained only with the simulated data. We utilize Matthews Correlation Coefficient (MCC) [47] as a balanced performance measure to compare inferred networks to the respective ground truth (see also [29]). Figures 2A and 2B show the results for the simulation study for all methods (see also Additional File 1, Tables S2, S3, S4 and S5). Overall, both *gLASSO_P_* and *BDgraph_P_* exhibit improved performance with relatively low standard deviation in terms of MCC as compared to their non-prior counterparts, both for low and high sample size settings. The performance of all other methods is affected by low sample sizes, with *BDgraph* showing slightly better performance than all other methods. Moreover, both *gLASSO_P_* and *BDgraph_P_* outperform all other methods as long as the prior noise does not exceed 10% *(gLASSO_P_*) and 30% of incorrect edges in the prior graph, in which case *BDgraph* achieves the highest median MCC over all methods. *GeneNet* performs well in all simulations, whereas *GENIE*3, *gLASSO* and *iRafNet* show about average performance with *iRafNet* achieving worst results overall. In addition to the curated prior matrices, we also generated a prior matrix reflecting the sparsity of the true graph (column ‘rbinom’ in Figure 2B and Additional File 1, Tables S2 and S3, see also Methods), and our results indicate, that information about sparsity of the underlying network already improves network inference performance. Finally, prior based methods, and specifically *BDgraph_p_*, outperform non-prior methods in the task of identifying the correct *cis* -gene by recovering associations between the discrete SNP and continuous gene expression data types (Additional File 1, Figure S3), when using independent eQTL data as prior.

**Figure 2:**
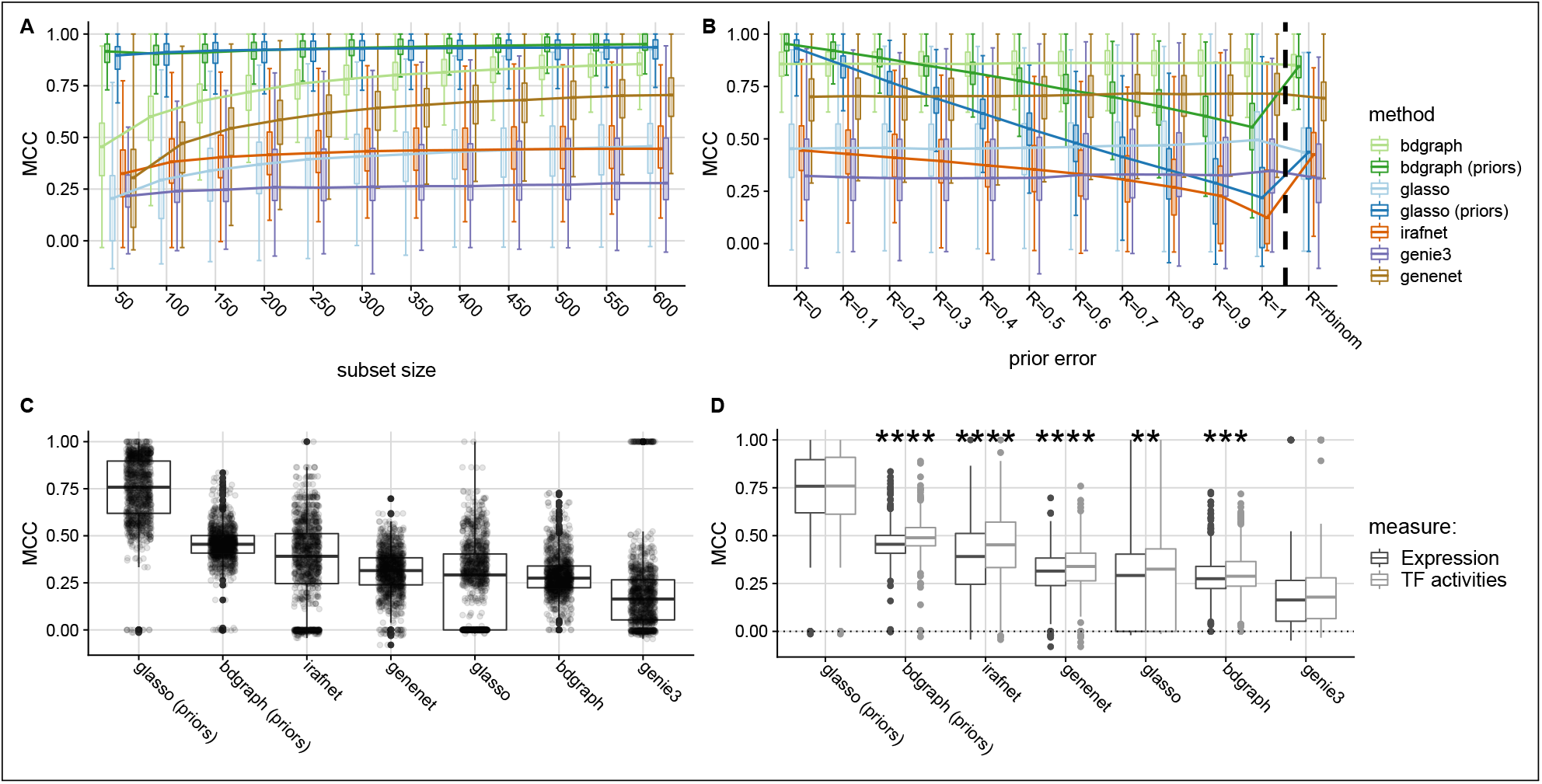
Method comparison results. **(A)** Results of simulation study: y-axis shows the Matthews correlation coefficient (MCC) as compared to the simulated ground truth, x-axis indicates increasing sample size from left to right, colors indicate different inference methods. **(B)** Similar to (A), but x-axis indicates increasing noise in the prior matrix from left to right. Group (‘*rbinom*’) indicates uniform prior set to reflect degree distribution of true graph. **(C)** shows MCC (y-axis) between networks inferred on KORA and LOLIPOP data for same locus for all methods (x-axis). **(D)** contrasts MCC across cohorts using TF expression (dark gray) versus using substituted TFAs (light gray). Boxplots show medians (horizontal line) and first and third quartiles (lower/upper box borders). Whiskers show **1**.**5** * *IQR* (inter-quartile range); for (**B**), dots depict individual results and for (**C**), stars indicate significant difference between expression/TFA results for each method (Wilcoxon test, **: *P* ≤ 0.01, ***: *P* ≤ 0.001, ****: *P* ≤ 0.0001)

#### Inferred networks replicate in independent datasets

In addition to the simulation study, we evaluated the methods on real world data from two large population cohorts: the KORA (Cooperative Health Research in the Region of Augsburg) and LOLIPOP (London Life Sciences Population) cohorts (see Figure 1A2 and Methods). Data from both cohorts were generated from whole-blood samples and contain imputed genotypes as well as microarray measurements of gene expression and DNA methylation for a total of 683 (KORA) and 612 (LOLIPOP) samples. Since for these data no ground truth is available, we evaluate robustness of the networks inferred by the individual methods via cross cohort replication. For each hotspot, we collect data for all genes, CpGs and the SNP in the locus set for KORA and LOLIPOP and separately inferred networks in both cohorts for all models. Obtained networks were then compared between cohorts using MCC to get a quantitative estimate of how robust the network inference is across different datasets for the same hotspot, yielding scores for KORA versus LOLIPOP and vice versa (i.e. one network functioning as the reference). Results of this analysis are shown in Figure 2C. With respect to MCC, models supplied with prior information (*gLASSO_P_*, *BDgraph_P_* and *iRafNet*) show the best performance, with *gLASSO_P_* coming up as the most robust method, followed by *BDgraph_P_* and *iRafNet*. Noticeably, of the top methods *BDgraph_P_* shows much less variance compared to *gLASSO_P_* and *iRafNet*. Ignoring prior information lead to a drop in performance for both *gLASSO* and *BDgraph*, which leads to *GeneNet* outperforming both methods. Finally, *GENIE*3 shows worst performance in this setting.

#### Estimated transcription factor activities as a proxy to TF activation

Transcription factor activities (TFAs) estimated from transcription factor binding sites (TFBS) and gene expression data have been suggested as an alternative to using TF gene expression in inference tasks [48], since a transcription factor’s expression level alone might not reflect the actual activity of a TF (driven for instance by its phosphorylation state). To evaluate, whether TFAs could improve our inference, we estimated TFAs for all TFs based on their expression and ChIP-seq derived TFBS from ReMap [49] and ENCODE [50, 51] (see Methods for details). We applied the same cross cohort replication strategy as above and compared MCCs from the TFA based analysis to the previous results using a one-sided Wilcoxon test. Figure 2D shows the results of TFA (light gray boxes) versus gene expression (dark gray boxes) based analysis in terms of MCC for all available hotspots. For all models but *gLASSO_P_* and *GENIE*3, TFAs yield a significantly higher MCC (Wilcoxon test *P* < 0.01) as compared to using the pure expression data (see also Additional File 1, Table S6).

According to the results presented above, detailed investigation of real world data was focused on networks obtained from *gLASSO_P_* and *BDgraph_P_* and TF expression was substituted by TFA estimates for all subsequent analyses.

### Replication of previous findings by simultaneous data integration

Before seeking new mechanistic insights and generating novel hypotheses from *trans* -QTL hotspots, we first checked whether our approach can replicate previous findings. Hawe *et al.* [10] inferred gene regulatory networks from *trans* -meQTL hotspots using a two-step approach involving 1) a random walk on a PPI and ChIP-seq based networks and 2) subsequent local correlation analysis. In contrast, our approach simultaneously integrates all functional data, relying on PPI and ChIP-seq information as prior knowledge, thereby avoiding the need for post-hoc correlation testing of e.g. SNP-gene and CpG-gene edges. For the comparison, we extracted three of their hotspot networks and evaluated the overlap with the networks inferred in this study.

**Table 2:** Comparison of the networks inferred in this study to the networks extracted from [10]. Numbers in bracket indicate statistics for the networks from the original publication.

| locus | num. nodes | num. edges | common edges | MCC |
| --- | --- | --- | --- | --- |
| rs9859077 | 99 (89) | 447 (287) | 141 | 0.52 |
| rs730775 | 58 (49) | 98 (67) | 48 | 0.69 |
| rs7783715 | 25 (17) | 24 (23) | 5 | 0.65 |

Table 2 shows the results of this comparison. Overall, the comparisons indicate relatively strong concordance between the two approaches with MCCs of 0.515, 0.689 and 0.65. Moreover, for all three networks, our simultaneous inference approach yielded more edges and nodes than the two-step approach (56%, 46% and 4% novel edges and 11%, 19%, 47% additional nodes for rs9859077, rs730775 and rs7783715, respectively), which might have been missed by the two-step approach, as it relies on known PPI and ChIP-seq information.

Figure 3 contrasts the two networks obtained for the *rs730775* hotspot using 1) the two-step approach by Hawe *et al.* [10] and 2) the network inferred in this study using *gLASSO_P_*, orange edges showing replicated and green edges indicating novel edges. In Hawe *et al.* [10], the authors described a regulatory network involving the *rs730775* SNP connected via *NFKBIE* to *NFKB1* which connects to the trans-CpG sites. This main pathway is also discovered in our approach (i.e. *rs730775 ↔ NFKBIE ↔ NFKB1 ↔ CpG sites*), in addition to some of the initially reported TFs (blue nodes), of which *NFKB1* is connected to most of the *trans* CpGs (82%, 29 out of 35) as was the case in the original network. However, we also identify patterns of CpG genes (green nodes) connected to the TFs, which were not previously identified. Overall, the integrated approach using prior information leads to high replication of previous networks including novel connections leading to potential new insights in target gene regulation.

**Figure 3:**
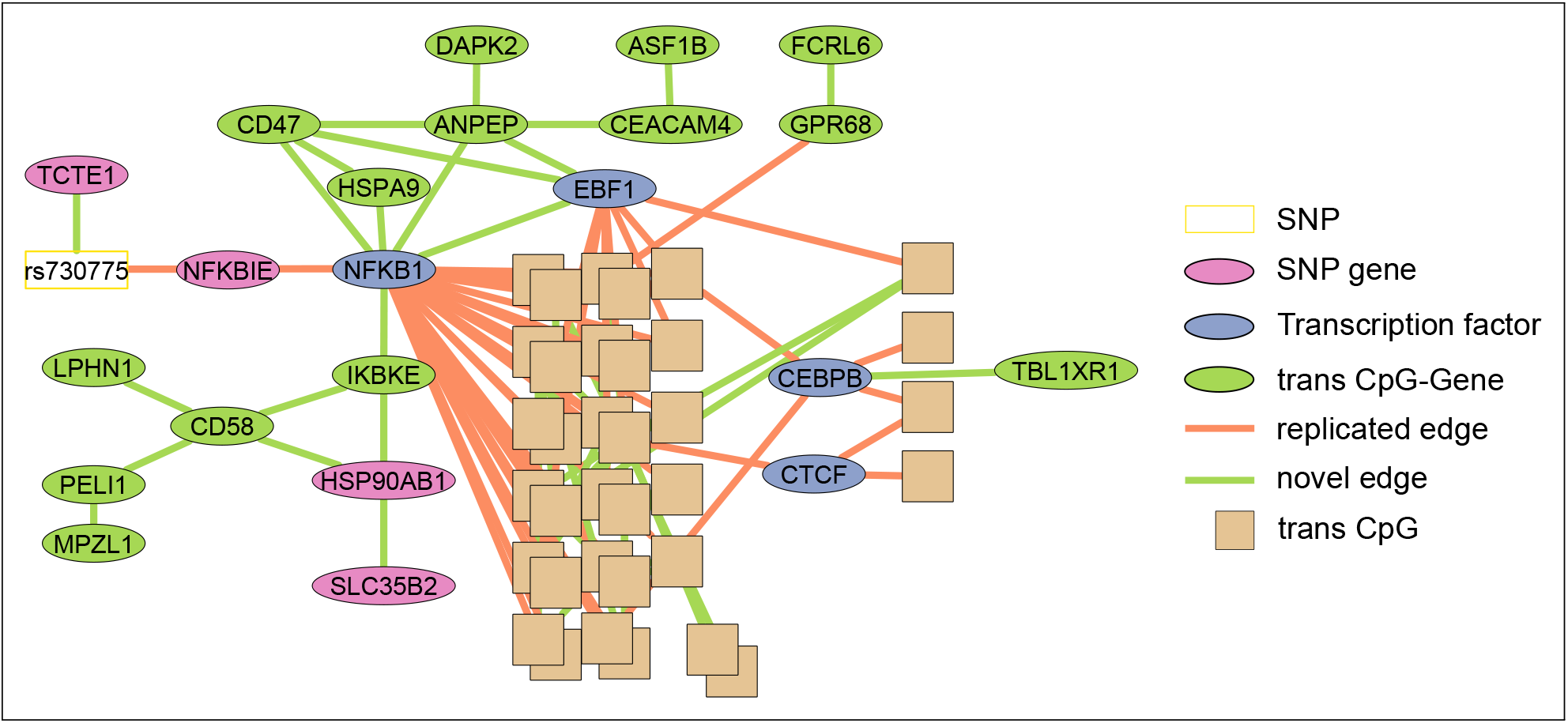
Comparison of the random walk based network reported in [10] and the network inferred from functional omics data in this study for the rs730775 locus. Shown is the complete network constructed from the omics data, edge color indicates replication/novelty. Orange edges: replicated with respect to the random walk network. Green edges: novel in our network. White box: SNP; pink nodes: SNP-genes; blue nodes: TFs; brown boxes: CpGS; green nodes: CpG-genes.

### A trans regulatory network for a schizophrenia susceptibility locus

In order to demonstrate the effectiveness of our approach in getting mechanistic insights from *trans* -QTL associations, we inferred networks for all meQTL [10] and eQTL [7] hotspots using whole blood data from the KORA and LOLIPOP cohorts using the prior based *gLASSO_P_* and *BDgraph_P_* models (see Methods, all networks are listed in Additional File 2, Table S3). Based on the GWAS catalog (v1.0.2, [52]), graph properties and a custom graph score (see Methods), we prioritized a *trans* acting locus that has previously been associated with schizophrenia (SCZ).

The network involves the *trans* -eQTL locus around the *rs9469210* (alias *rs9274623*^1^ SNP in the Human Leukocyte Antigen (HLA) region on chromosome 6 shown in Figure 4A.

**Figure 4:**
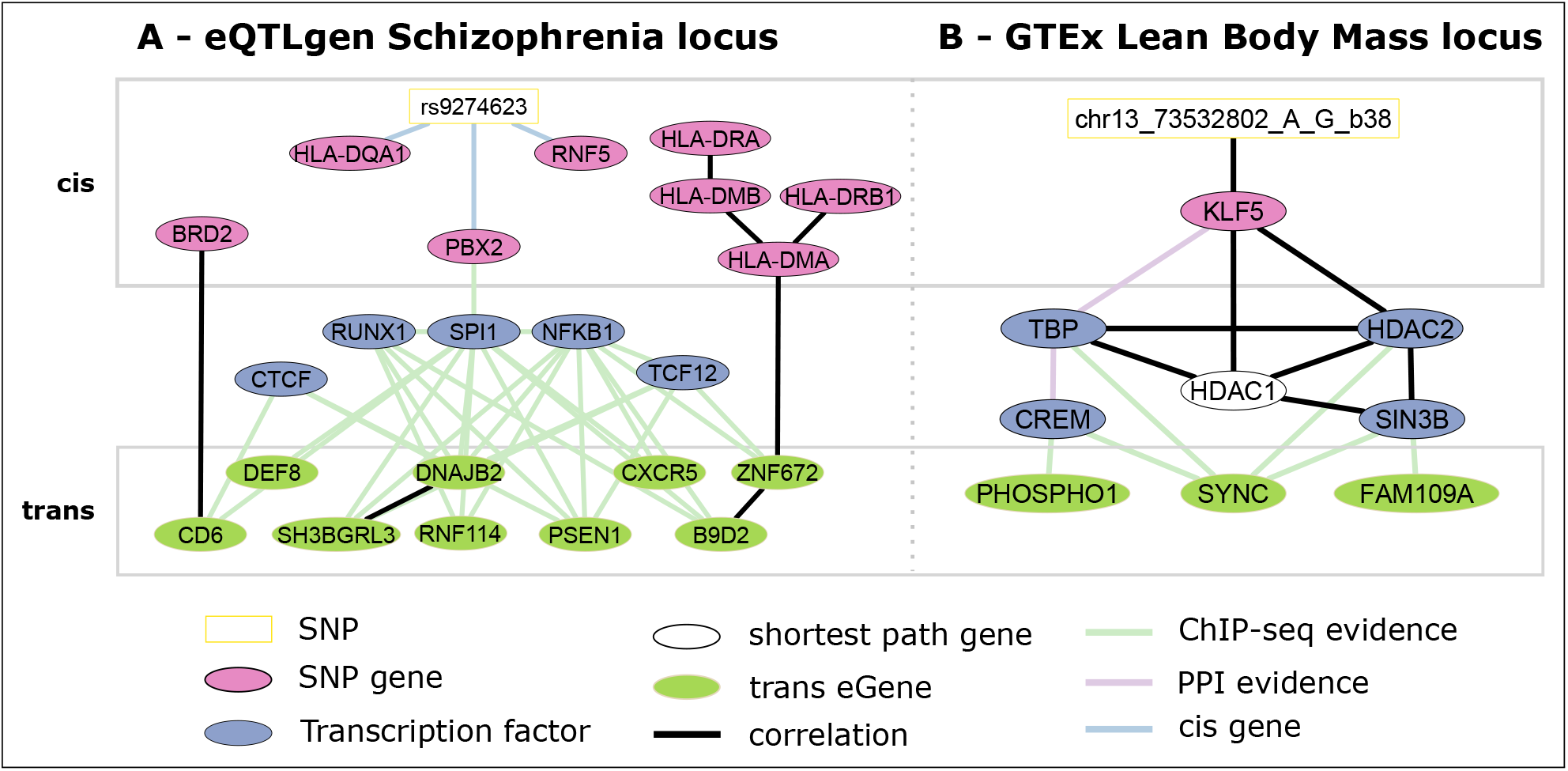
Inferred networks for the schizophrenia susceptibility locus rs9274623 obtained from eQTLgen (A) and the rs9318186 locus obtained from GTEx (B). The white boxes indicate sentinel SNPs, pink ovals indicate SNP-Genes, blue ovals transcription factors and white ones shortest path derived genes. Light green ovals represent genes trans-associated to the SNP. Black edges were inferred during network inference. In addition to being inferred, colored edges indicate ChIP-seq protein-DNA binding evidence (green), protein-protein interaction in the BioGrid (purple) and whether or not a gene is encoded in *cis* of the linked entity (blue).

*rs9274623* has been associated with SCZ [54] and is a *cis* -eQTL for all three of its directly connected SNP-genes, *PBX2, RNF5* and *HLA-DQA1* in the eQTLGen study. *RNF5* showed differential expression for SCZ cases vs controls in addition to its expression being associated with an additional independent SCZ susceptibility SNP (rs3132947, *R*^2^ = 0.14 in 1000 genomes Europeans^2^) located in the HLA locus [55]. Interestingly, *PBX2* has been associated with a SCZ related phenotype in a pharmacogenetics study (clozapine-induced agranulocytosis) [56, 57] and shows direct binding evidence to the *SPI1* promoter region (ReMap TFBS [49]). The transcription factor *SPI1 (PU.1*) is linked to Alzheimer’s Disease likely by impacting neuroinflammatory response [58] and was found to interact with its network neighbor,*RUNX1*, in modulating gene expression [59]. Moreover, *RUNX1* has been implicated in rheumatoid arthritis, a disease negatively associated with SCZ and which hence might share susceptibility genes with SCZ [60]. Interestingly, several genes encoded in the HLA locus, which has been implicated in SCZ and other psychiatric and neurological disorders [61–64], were picked up by our inference downstream of *SPI1* and *RUNX1. TCF12* is a paralog of *TCF4* and *TCF3* which are known E-box transcription factors and are expressed in multiple brain regions [65]. *TCF4* loss-of-function mutations are the cause of Pitt-Hopkins syndrome (a syndrome causing mental retardation and behavioral changes amongst other symptoms) [66] and regulatory SNPs relating to *TCF4* have been associated with SCZ [67, 68]. The *NFKB1* pathway has been recognized as an important regulatory and developmental factor of neural processes and was found to be dysregulated in patients with SCZ [69]. Finally, 9 of the 40 discovered *trans* -eGenes of the locus are connected to the SNP via the selected TFs. Of these, *SH3BGRL3* [70] has already been linked to SCZ and *PSEN1* [71], *B9D2* [72], *CXCR5* [73] as well as *DNAJB2* [74] were implicated in other neurological disorders. In addition, the *trans* eGene *RNF114* has previously been shown to play a role in the *NFKB1* pathway [75]. A formal colocalization analysis using fastENLOC [76] showed evidence of a common causal variant underlying the SCZ GWAS signal [77] and each of the eQTLGen *trans* -eQTL of PSEN1, DNAJB2 and CD6 (SNP-level colocalization probability of 0.92, 0.87 and 0.42, respectively; see Methods and Additional File 1, Figure S4).

Our approach highlighted a potential regulatory pathway involving diverse genes related to SCZ and other neurological disorders. While some of the genes were not previously reported in this specific disease context (e.g. *CD6, BRD2, DEF8*), their association to this network indicates a potential role in SCZ pathogenesis and additional colocalization analysis hints at a potential causal relationship between these genes and SCZ.

### Application to GTEx Skeletal Muscle tissue

All above analyses were focused on whole-blood data, however, the proposed strategy can be applied to data from any biological context. To demonstrate this, we investigated the recently published *trans* -eQTLs from the GTEx v8 release [38, 78]^3^. We identified a single LD block in Skeletal Muscle tissue, which is a *trans* -eQTL hotspot (see Methods), and for which we inferred regulatory networks. Since we can’t use the same priors, which were initially derived from GTEx, to analyze the same data set, we set out to curate muscle tissue specific priors from independent datasets. We utilized muscle eQTL from Scott *et al.* (2016) [79] and gene expression data curated from the ARCHS^4^ [41] database and generated tissue specific TFBS using factorNet [80] on DNAse-seq data obtained from ENCODE [50, 51]4 (see Methods for details). The resulting network for the *gLASSO_P_* model is shown in Figure 4B.

The genetic variant rs9318186 is a *cis* -eQTL of *KLF5* in GTEx v8 Skeletal Muscle (*P* = 6.1*x*10^-37^) and a proxy of it (*R*^2^ = 0.88) has been associated with *Lean Body Mass* (LBM). *KLF5* itself, too, has been associated with LBM in a transcriptome-wide association study integrating GWAS results with gene expression [81] and with lipid metabolism in *KLF5* knockout mice [82]. In addition, several other genes in the network have been associated with related phenotypes: Both *HDAC1* and *HDAC2* have been found to control skeletal muscle homeostasis in mice [83], work together with *SIN3B* in the SIN3 core complex to regulate gene expression and are involved in muscle development [84]. TATA binding protein (*TBP*) is a well known transcription factor and important for the transcriptional regulation of many eukaryotic genes [85]. The *trans* -eGene *SYNC* was found to interact with dystrobrevin (*DMD* gene) in order to maintain muscle function (during contraction) in mice as well as being associated with neuromuscular disease [86, 87]. In addition, in Seim *et al.* (2018) [88], the authors investigated the relationship between obesity and cancer subtypes and found, that both *PHETA1* /*FAM109A* expression are associated to Body-Mass-Index (BMI) in esophageal carcinoma in data from The Cancer Genome Atlas (TCGA). *PHOSPHO1* has been found to be involved in metabolism, specifically in energy homeostasis [89], and has also been associated via DNA methylation with BMI [90, 91] and with HDL levels, which have been negatively associated with LBM [92]. Dayeh *et al.* (2016) [93] further showed decreased DNA methylation at the *PHOSPHO1* locus in skeletal muscle of diabetic vs. non-diabetic samples. The remaining gene in the network *(CREM*) has not yet been described in the broader context of LBM, but a GWAS meta-analysis executed by Wang *et al.* (2014) [94] hinted at association of a *CREM* SNP (rs1531550, *P* = 1.88×10^-6^) with elite sprinter status. These results suggest, that *KLF5* may exert its specific functions through transcriptional regulation via the SIN3 core complex including *TBP*, with a potential involvement of *CREM*, of the *trans* -eGenes *PHOSPHO1, SYNC* and *PHETA1/FAM109A.*

## Discussion

In this study, we introduced a Bayesian framework for the inference of undirected regulatory networks underlying molecular *trans* -QTL hotspots across multi-omics data types using existing prior knowledge. We compiled a comprehensive set of context specific network edge priors from diverse biological databases and applied these together with multi-omics data in different settings. These settings include an extensive simulation study to benchmark state-of-the-art inference methods as well as application to two large population cohorts, which we use for a replication analysis on the one hand and to generate novel hypotheses about molecular disease mechanisms on the other hand. Moreover, by applying our approach a GTEx Skeletal Muscle eQTL hotspot, we showed, that our strategy can be applied to data sets from other tissues, generated with different technologies.

Benchmarking is important for selecting the best possible methods for specific tasks and we hence followed recently published guidelines [95] to perform benchmarking of state-of-the-art network inference methods in 1) a simulation study and 2) a replication analysis. Results from both analyses were then used to select the methods best suited for network inference based on functional multi-omics data from QTL hotspots using prior information.

By inferring networks in over 10,000 simulated data sets, which reflect the distribution of network parameters obtained from real-world data, we showed, that methods utilizing prior information outperform methods without any prior information in recovering a simulated ground truth, similar to what has been found e.g. in [27, 28, 36]. We further observed that, as expected, too much noise in the prior information significantly reduces method performance. However, only by increasing the noise level, i.e. the percentage of incorrect prior edges, to above 30% decreases the performance for BDgraph below the performance of its non-prior counterpart, indicating that low levels of noise in edge priors still improve network inference, results which are in line with e.g. Wang *et al.* (2013) [30], who used a modified graphical lasso approach, Christley *et al.* (2009) [28], who used an regularized ODE model and Greenfield *et al.* (2013) [27], who used a Bayesian regression framework. We further find, that, both for the prior and non-prior case, the Markov-Chain-Monte-Carlo based *BDgraph_P_* method outperforms respective other methods. However, both the copula approach based BDgraph and the *gLASSO_p_* outperform other methods in recovering mixed edges between discrete SNP allele dosage and continuous gene expression levels, although the tree based methods should be able to incorporate mixed data. While *BDgraph_P_* shows overall better performance than *gLASSO_p_*, the graphical lasso exhibits much lower run time which can be an important practical consideration. Our results hence highlight the strong value of using prior information for multi-omics based network reconstruction, and slightly favor BDgraph over the graphical lasso for this kind of inference.

We confirmed the results of the simulation study by extended benchmarking of inference methods in a cross cohort replication analysis on two large multi-omics data sets. Prior based methods showed overall best replication across different cohorts as compared to non-prior methods. In the real-world setting, however, *iRafNet* performed similarly well as the other two prior methods in contrast to the simulation study and all prior based methods outperform non-prior methods. The good replication of prior based methods across different cohorts shows, that curated priors help to obtain more stable and confident results as compared to using functional data alone. Together with the simulation, these results provide a comprehensive benchmark of established network inference methods and suggest, that priors should be integrated in network inference tasks wherever possible.

Based on the results from the replication and simulation study, we choose the two best (prior based) methods *BDgraph_P_* and *gLASSO_P_* for detailed investigation of networks obtained form real-world cohort data. Using our integrative approach, we were able to reproduce and expand upon previous results from a step-wise network analysis approach presented in [10]. Of three of the locus networks described in their study, we reconstructed most of the edges and found additional edges, allowing more mechanistic interpretations for the function of specific transcription factors in relation to DNA methylation. One reason for finding additional edges is, that these could not be detected by the previous approach, since the authors focused on using established PPI and protein-DNA interactions and did not test all possible edges in the functional data. In contrast, our integrated approach considers all edges regardless of available prior evidence and associations will emerge, if the signal in the functional data alone or in addition to the prior evidence is strong enough.

Next, we utilized the two top performing methods (*BDgraph_P_* and *gLASSO_P_*) to infer networks from *trans* -eQTL hotspots and found, that our strategy can be used to recover known biology on the one hand and generate novel hypotheses about the molecular basis of diseases on the other hand. For a schizophrenia (SCZ) susceptibility locus, we identified several known SCZ (e.g. *RNF5, HLA genes* [55, 61]) and related (e.g. *PBX2* [56, 57]) genes in the inferred locus network. Caution is needed for the interpretation of the candidates based on *cis* -eQTL, because of the haplotype structure of the HLA locus. However, our candidate *PBX2* is defined by its connections in the network to the *trans* genes and, therefore, independent of the *cis* eQTL. Expanding upon similar previous observations based on *trans* eQTL [7], the integrated network analysis including associated *trans* genes prioritizes *PBX2*, which was not possible using *cis* -eQTL alone. It was previously hypothesized, that *RUNX1* is involved in SCZ due to a negative association of SCZ with rheumatoid arthritis [60]. Our network corroborates this hypothesis and further allows for generating novel hypotheses about the involvement of other genes (e.g. *BRD2, DEF8* and *RNF114*), which could potentially play a role in schizophrenia. Moreover, we further substantiated these results by a formal colocalization analysis of the *trans* -eQTL and schizophrenia GWAS [77] signals of the *trans* genes linked in the network, which revealed strong evidence for colocalization of the underlying genetic variants of the disease and molecular traits. As this locus was derived from whole-blood data, interpretation is not straight forward for SCZ. Ideally, this analysis can be followed up in data derived from brain tissue to corroborate findings.

To show, that our approach can be applied across different omics types and data sets, we analyzed a Skeletal Muscle *trans* -eQTL hotspot from GTEx associated with Lean Body Mass. We recovered known genes involved in lipid metabolism *(KLF5* [81, 82]) as well as muscle development and controlling skeletal muscle homeostasis (e.g. *HDAC1*, *HDAC2*, [83]) and maintaining muscle function (*SYNC* [87]). This shows, that the genes linked in the inferred network are overall coherent with the observed phenotype association at this *trans* -acting locus. Moreover, *HDAC1*, *HDAC2* and *SIN3B* have been described to interact together during muscle development [84], and, although these results were described in mice, our results suggest that these genes could exhibit a similar function in human. In addition, we observed an association between *CREM* and *SYNC* in our network, which led us to hypothesize, that *CREM* might also be involved in maintaining muscle function and Lean Body Mass, although is has not been previously linked to these phenotypes. However, additional experimental validation needs to be performed in order to corroborate findings of these computational analyses.

Several practical considerations arise from our findings: First, by investigating the effect of increasing amounts of noise in the prior information in our simulation study, we showed, that some caution needs to be applied when curating continuous prior information from public biological data to keep noise levels low. Therefore, although *gLASSO_P_* and especially *BDgraph_P_* seem to be robust to low to moderate levels of noise, one might consider using only experimentally validated protein-protein interactions or high quality gene expression data to generate priors. Next, the definition of hotspot locus sets and priors in this study mitigates the *N << P* problem. This has been a problem sought to be alleviated using specialized approaches in previous applications [4]. Using our approach, the total number of entities (variables) going into the network inference typically does not exceed the total number of available samples in our data sets, and we showed in a simulation study, that priors improve inference also in low sample size settings. Overall, the benefit of the locus sets comes with the risk of missing certain genes needed to fully describe the *trans* effects. For instance, we reason that most relevant genes lie on the shortest path between *cis* and *trans* entities in the PPI network and hence only included those shortest path genes. However, our strategy of curating a stringent set of relevant transcription factors as well as including genes showing protein-protein interactions and all the genes in the vicinity of the hotspot SNP, should enable most key regulator genes to enter the inference process and yields parsimonious and easily interpretable results. In addition, methods have been developed to handle mixed data types, such as e.g. genotypes and gene expression. *BDgraph*, which uses a copula based approach to transform non-normal data, showed better performance in recovering associations between discrete and continuous data types as compared to *gLASSO* and the tree based methods, and hence should be preferred for applications on mixed data, especially when prior information is available. Finally, while we could use transcription factor binding sites (TFBS) in blood related cell-lines to analyze whole-blood cohort data, context (e.g. tissue) specific TFBS are not yet available for a large number of transcription factors, which potentially limits this approach to fewer applications. However, novel developments to predict TFBS from context specific open chromatin information (e.g. *factorNet* [80]) can help in carrying this strategy to more contexts. As an example, we utilized TFBS predicted using *factorNet* based on ENCODE [50, 51] DNAse-seq data for analyzing a GTEx Skeletal Muscle *trans* eQTL locus.

## Conclusion

This study describes a novel strategy for using comprehensive edge-wise priors from biological data to improve network inference for *trans* -QTL hotspots from human population scale multi-omics data. This facilitates the investigation of their underlying regulatory networks and enables the generation of novel mechanistic hypotheses for disease associated genetic loci. Moreover, we report a rigorous benchmark of state-of-the-art network inference methods for this task both in simulated and real-world data, and highlight the benefit of including biological prior information to guide network inference.

## Methods

### Cohort data processing

Methylation data were measured using the Infinium Human Methylation 450K BeadChip in both the KORA and the LOLIPOP cohort and methylation beta values obtained as described previously [43, 44]. Quantile normalized methylation beta values were adjusted for Houseman blood cell-type proportion estimates and the first 20 principal components calculated on the array control probes by using residuals of the following linear model:

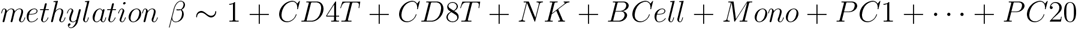

For expression data, the Illumina HumanHT-12 v3 and Illumina HumanHT-12 v4 expression BeadChips were used in KORA and LOLIPOP, respectively, and processed as described previously [10, 96]. Only probes common to both arrays were selected for analysis. Expression data were adjusted for potential confounders by regressing log2 transformed expression values against age, sex, RNA integrity number (RIN) as well as RNA amplification plate (KORA) / RNA conversion batch (LOLIPOP) (batch1) and sample storage time (KORA) / RNA extraction batch (LOLIPOP) (batch2) and obtaining the residuals from the linear model:

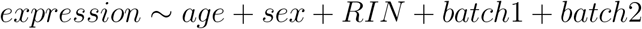

Additional details on the cohort data and design are presented in [43, 96, 97] (KORA) and [44, 98] (LOLIPOP).

For the inference of the GTEx Skeletal Muscle related network, we used GTEx v8 Skeletal Muscle data [78]. Potential confounders including first 5 genotype PCs, 60 expression PEER factors and measured covariates ‘WGS sequencing platform’ (HiSeq 2000 or HiSeq X), ‘WGS library construction protocol’ (PCR-based or PCR-free) and donor sex, were removed from expression data prior to analysis. Processing has been performed as previously described and details can be found elsewhere [78].

### Hotspot extraction and construction of locus sets

We extract sub-sets of genomic entities (SNPs, CpGs and genes) on which we perform network inference based on the *trans* -meQTL reported by [10] (Supplementary Table 9 of their study) and eQTLGen *trans* -eQTL [7]^5^. For GTEx, we obtained current (GTEx v8) tissue specific *trans* -eQTL from https://www.gtexportal.org/home/datasets^6^.

### Hotspot extraction

The list of *trans* -meQTL results obtained from [10] was already pruned for independent genetic loci and was used as provided in the paper supplement. To remove redundant highly correlated genetic loci, we pruned the eQTLGen *trans* -eQTL by selecting the eQTLs with 1) the highest minor allele frequency and 2) the largest number of trans genes for each LD cluster (1Mbp window, *R*^2^ > 0.2). For GTEx, we merged eQTL by combining SNPs with *R*^2^ > 0.2 and distance < 1Mbp to independent genetic loci and kept all *trans* -eGenes (eGenes: genes associated with eQTL genotype) of the individual SNPs for this locus. The SNP with the highest MAF was selected as a representative SNP for the hotspot. We defined hotspots as genetic loci with ≥ 5 *trans* associations, yielding a single hotspot for GTEx, 107 for the meQTL and 444 for the eQTLGen data (Additional File 2, Tables S1 and S2). In [10], the authors provide a total of 114 meQTL hotspots per our definition. We discarded 7 of the 114 meQTL hotspots (SNPs rs10870226, rs1570038, rs17420384, rs2295981, rs2685252, rs57743634, rs7924137, as either no *cis* genes are available or no gene expression data were measured for any of the annotated *cis* genes (mostly lincRNAs, miRNAs and pseudogenes; Additional File 1, Table S1), which are needed for locus set definition (see below).

### Locus sets

To mitigate the *N << P* problem in network inference [4], where the number of features or parameters far exceeds the number of samples, we run the inference on a subset of genomic entities (SNPs, genes and CpGs) induced by *trans* hotspots. We therefore gathered all genes, which could be involved in mediating the observed QTL effects and thus were considered during the network inference, in the form of *locus sets* for each hotspot. We bridge the gap between the involved chromosomes by including transcription factor binding site (TFBS) information collected from *ReMap* [49]^7^ and *ENCODE* [50, 51]^8^ as well as human protein-protein interaction (PPI) information available via *theBioGrid* [99]^9^ (version 3.5.166). We filtered *ReMap* and *ENCODE* TFBS for blood related cell types by selecting all samples which contain at least one of the following terms: “amlpz12_leukemic”, “aplpz74_leukemia”, “bcell”, “bjab”, “bl41”, “blood”, “lcl”, “erythroid”, “gm”, “hbp”, “k562”, “kasumi”, “lymphoblastoid”, “mm1s”, “p493”, “plasma”, “sem”, “thp1”, “u937”. Genes in the PPI network were filtered for genes expressed in whole blood (GTEx v6p *RPKM* > 0.1)^10^. We enumerated all entities to be included in the locus set by performing the following steps:

1. Define set *S_L_* for a locus *L* and add the QTL entities (QTL SNP 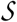 and *trans* -QTL eGenes/CpGs 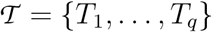, where *q* is the number of associated *trans* entities for *L*)
2. Add all genes encoded within 500kb (1Mbp window) of 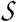 as **SNP-Genes** to *S_L_* (set 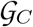)
3. For meQTL hotspots, add genes in the vicinity of each 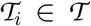 (previous, next and overlapping genes with respect to the location of 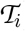) as **CpG-Genes** to *S_L_* (set 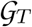)
4. Add all **TFs** with binding sites within 50bp of each CpG or binding in the promoter region of each gene over all 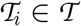 to *S_L_* (set 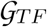)
5. Add shortest path genes *G_SP_*, i.e. genes which connect 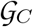 (step 2) with 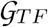 (step 4) according to BioGrid PPIs to *S_L_*

To define *G_SP_*, we added only genes which reside on the shortest path between the *trans* entities 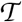 and the SNP-Genes 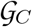 in the induced PPI sub-network, i.e. containing all genes and their connections which can be linked to either 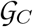 or the TFs 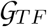. Specifically, we added the CpGs to the filtered BioGrid PPI network, connected them to the TFs 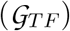 which show binding sites in their vicinity and calculated node weights based on network propagation as described in [10]. We then extracted nodes on paths with maximal total propagation score based on node-wise propagation scores *PS*. For this, we weighted node scores proportional to (−1) × *PS* and then calculate the minimal node-weight paths between *trans* entities 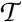 and SNP-Genes 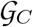 using the *sp.between()* method of the *RBGL* R package (version 1.56.0, R interface to the Boost Graph Library [100]) and extracted all genes on the resulting shortest paths. All nodes of the generated locus set were subsequently used as inputs to the network inference.

### Prior generation

We utilized several data sources to define priors for possible edges between and within different omics levels. Each possible edge between entities in the locus set can only be assigned a single type of prior. Specifically, the different priors include:

- **SNP-to-Gene** priors, for edges between the SNP 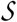 and SNP-Genes 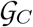
- **Gene-to-Gene** priors, for edges between all gene-gene combinations except TFs 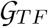 and their eQTL based targets in 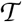
- **CpG-to-Gene** priors, for edges between CpGs in 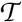 and their neighbouring genes 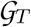
- **TF-to-target** priors, for edges between TFs 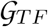 and their targets in the *trans* set 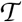

#### SNP-to-Gene

To obtain SNP-to-Gene edge priors, we downloaded the full GTEx v6p whole-blood eQTL table^11^) and calculated, for each SNP-Gene pair, the local false discovery rate (lFDR, [101]) using the *fdrtool* R package (version 1.2.15). As described in Efron *et al.* (2008) [101], the lFDR represents the Bayesian posterior probability of having a null case (i.e. that the null hypothesis is true) given a test statistic. We therefore defined the prior for a specific SNP 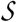 and a SNP-Gene 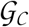 as 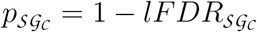.

#### Gene-to-Gene

We formulate *Gene-to-Gene* edge priors by combining public GTEx gene expression data [38] with PPI information from the BioGrid [99] to retrieve co-expression p-values and the respective lFDR for pairs of genes connected by a protein - protein interaction. A special case are priors between TFs and their target genes as identified via ChIP-seq (see above), which are not considered as *Gene-to-Gene* edges but are handled separately as described under ‘TF-to-target priors’ below. GTEx v6p RNA-seq gene expression data were downloaded from the GTEx data portal^12^. Expression data for GTEx were filtered for high quality samples (RIN ≥ 6) and log2 transformed, quantile normalized and transferred to standard normal distribution before removing the first 10 principle components to remove potential confounding effects [102]. Priors were derived for all Gene-Gene pairs with PPIs in the BioGRID network, where a gene 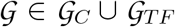 (for meQTL) or 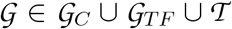 (for eQTL). For each pair, we calculated the Pearson correlation p-values in the GTEx expression data and subsequently determined the lFDR over all p-values. The prior for two genes 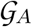 and 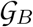 was then set to 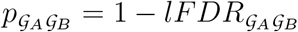.

#### CpG-to-Gene

For the *CpG-to-Gene* priors (meQTL context only), we utilized two strategies, distinguishing between TF-CpG priors (i.e. priors between CpGs and TFs showing binding sites near the CpG site, described below under ‘TF-to-target priors’) and CpG-to-Gene priors (i.e. where the gene itself is encoded near the CpG). For the *CpG-to-Gene* priors, we utilized the genome-wide chromHMM [103] states (15 states model) identified in the Roadmap Epigenomics project [40]^13^. These states reflect functional chromatin states in 200bp windows and were obtained using histone mark combinations as identified via ChIP-sequencing. We quantified a CpGs potential to affect a nearby gene, *pT_x_*, by retrieving the proportion of Roadmap cell-lines in which the CpG resides within a transcription start site (TSS) related state (see Table 3). We further adjusted the *pT_x_* by weighting state information according to the Houseman blood cell type estimates available from our data. To this end, we took the population mean for each of the Houseman cell proportion estimates and multiplied them with the chromHMM state proportions. A specific CpG-to-Gene prior for a CpG 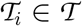 and a gene 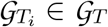 was then set to 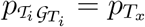, if the genomic distance 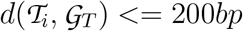.

**Table 3:** Description of chromHMM states used in our analyses as given at https://egg2.wustl.edu/roadmap/web_portal/chr_state_learning.html. Bold faced states were defined as ‘active transcription’ states and used to set CpG-Gene priors.

| STATE NO. | MNEMONIC | DESCRIPTION |
| --- | --- | --- |
| **1** | **TssA** | **Active TSS** |
| **2** | **TssAFlnk** | **Flanking Active TSS** |
| 3 | TxFlnk | Transcr. at gene 5' and 3' |
| 4 | Tx | Strong transcription |
| 5 | TxWk | Weak transcription |
| 6 | EnhG | Genic enhancers |
| 7 | Enh | Enhancers |
| 8 | ZNF/Rpts | ZNF genes & repeats |
| 9 | Het | Heterochromatin |
| **10** | **TssBiv** | **Bivalent/Poised TSS** |
| **11** | **BivFlnk** | **Flanking Bivalent TSS/Enh** |
| 12 | EnhBiv | Bivalent Enhancer |
| 13 | ReprPC | Repressed PolyComb |
| 14 | ReprPCWk | Weak Repressed PolyComb |
| 15 | Quies | Quiescent/Low |

#### TF-to-target priors

We formulate separate priors for all edges between transcription factors 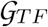 and *trans* CpGs (meQTL) and *trans* genes (eQTL) in 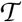. Priors were only set for TF-to-CpG edges were we observe a TF binding site (from ReMap/ENCODE, see above) within 50bp of the CpG. For TF-to-Gene edges, we only considered pairs were the TF has a binding site 2,000bp upstream and 1,000 downstream of the gene’s TSS. In both cases, if the TFBS criteria are met, we set a fixed large prior of 0.99 for all 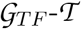 pairs to represent the strong protein-DNA interaction evidence of ChIP-seq data.

Finally, the priors for all remaining possible edges which were not set based on one of the criteria described above, e.g. for SNP-to-Gene edges without eQTL in the GTEx data, were set to a small pseudo-prior *p_pseudo_* = 10*e*^-7^.

### Ground truth network generation, data simulation and prior randomization

We performed a simulation experiment for each of the meQTL hotspots. For each SNP 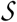 and its corresponding locus set 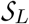, we first collect the corresponding prior matrix 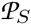 with priors defined as described above. We generate 10 noisy 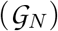 ground truth graphs 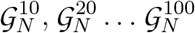 by switching edges in the graph while keeping the degree distribution of a sampled graph 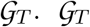 is generated using all entities of 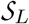 by uniformly sampling from 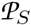, i.e. 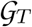 contains an edge *e_ij_* for each element *p_ij_* of 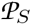, if *p_ij_* > *p_pseudo_* and *runif* (0,1) = *p_ij_*, where *runif*(0,1) generates uniformly distributed random numbers between [0,1]. This procedure effectively introduces noise in the study. For instance, by switching 10% of the edges from 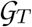 to generate 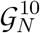, and making sure, that the new edges are not present as priors in 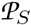, we introduce a noise level of 10% when comparing 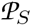 to 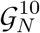. We simulate data for each 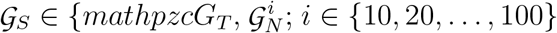 using the *bdgraph.sim()* method of the *BDgraph* package with parameters: 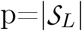 (number of nodes), 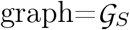, N=612 (number of samples in LOLIPOP) and mean = 0. This approach generates normally distributed data with a covariance structure as defined by the ground truth graph. We want to assess the impact of having discrete (genotype) data present for the network inference. To this end, we converted the SNP variable in the simulated data to genotype dosages (0,1,2) reflecting the allele frequencies of the genetic variant used in this simulation run. Specifically, we transformed the Gaussian data obtained from *bdgraph.sim()* to discrete values using the frequencies of the individual dosages for the SNP in the LOLIPOP data as quantile cut points. For each of these simulated data individually, we infer the network models and compare the inferred networks to the respective ground truth graphs 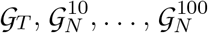. We added one additional comparison, evaluating a prior on the density of the observed graph. For this, we estimated a single prior value reflecting the desired density for all edges based on a binomial model. We use the number of edges 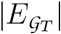 of all sampled graphs 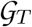 for a single run, the total number of possible edges |*E_T_*| = (*N* * (*N* – 1))/2, with *N* the total number of available nodes, and set the prior as

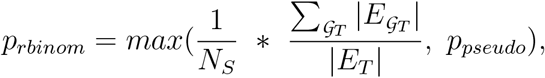

where *N_S_* is the number of sampled graphs (i.e. the number of randomizations). For each hotspot, we repeated the above simulation procedure 100 times to obtain stable results.

### Network inference

Based on the data and priors gathered for the individual hotspots, we set out to infer the regulatory networks which are best supported by these data. We evaluated several state-of-the art methods with respect to their applicability to this problem, both in a simulation study (see above) and via replication of inferred networks in real-world data from two large human population based cohorts. We applied *GeneNet* [45, 104], the graphical lasso [*glasso*, 42], *BDgraph* [29], *iRafNet* [32] as well as *GENIE3* [46] on the individual data to reconstruct regulatory networks using the respective *CRAN*^14^ and *bioconductor*^15^ R packages. An overview on the used inference methods and package versions is given in Table 1. Methods were chosen to reflect a range of different approaches (i.e. shrinkage based partial correlation in *GeneNet*, Bayesian MCMC sampling in *BDgraph*, lasso in *gLASSO* and tree based inference in *iRafNet* and *GENIE***3**), based on whether or not implementation was readily available and whether prior knowledge could be incorporated. The well known *GeneNet* and *GENIE***3** methods are not capable of utilizing prior information, but were used as a reference for comparison to the other methods.

#### GeneNet

For the application of GeneNet we first filtered any CpG probes from the data containing missing values. We then estimated the regulatory network by calling first the *ggm.estimate.pcor* followed by the *network.test.edges* and *extract.network* methods, all with default parameters.

#### GENIE3

To infer networks with GENIE3, we again used the NA filtered data (see above) with the *GENIE3* method of the package followed by the *getLinkList* method using default parameters. GENIE3 generates a ranked list of regulatory links which do not relate to any statistical measure and hence a cutoff for the link weights has to be identified manually^16^. To define an optimal cutoff, we first divide the list of weights into 200 quantiles (marking 200 distinct cutoffs) if the number of unique link weights exceeded 200. We then extracted for each cutoff the respective regulatory network and compared it to a scale free topology analogously to the approach used in [105], generating *R*^2^ values indicating the goodness-of-fit to the topology. To choose the final network, we followed the approach suggested by Zhang *et al.* (2005) [105], which suggests to use networks with *R*^2^ > 0.8. If none of our networks fit that criteria, we choose the network with the highest *R*^2^.

#### BDgraph

We used BDgraph to infer networks under consideration of prior information as well as without prior information (*BDgraph* and *BDgraph_P_*) using the *bdgraph* method of the *BDgraph* CRAN package (version 2.61). The following parameters were set: *method = “gcgm”, iter = 10000, burnin = 5000.* We further set the *g.prior* parameter to the prior matrix collected for the hotspots and the *g.start* parameter to the incidence matrix obtained from the prior matrix by setting all entries with prior information > 0.5 to 1 and all others to 0. For comparison with the no prior case, we kept all parameters the same but omitted the *g.start* and *g.prior* parameters. The graph was then obtained from the fitted model using the *select* method of the package with parameter *cut = 0.9,* thereby only choosing edges with a posterior probability of at least 0.9.

#### glasso

Similar to BDgraph, we utilized the graphical lasso both with and without prior information. To infer the graphical lasso models, we used the *glasso* method available in the *glasso* CRAN package and set the parameter *penalize.diagonal = FALSE.* The *glasso* takes a regularization parameter λ, which implies either strong penalization of edges (high λ) or weak penalization (low λ) of parameters. This parameter can also be supplied as a matrix Λ of size *n × n* (where *n* is the number of nodes/variables) in order to supply individual parameters for individual edges. We integrated the prior information by first transforming the prior matrix 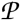 such that 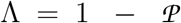 and then supplying Λ as the regularization matrix containing values for each possible edge. This approach is similar to what has been proposed in [30, 31]. In addition, we screened a selection of penalization factors *ω* for both the prior and the none prior case to construct the optimal graphical lasso network with respect to the Bayesian Information Criterion (BIC). For the prior case, we included *ω* in the model by setting Λ = Λ × *ω*). For the non-prior case, we set *λ* = *ω*. We performed 5-fold cross validation and inferred the model for all *ω ∈* {0.01,0.015,..., 1} on the training set (containing 80% of the data) and then selected the *ω* yielding the minimal mean BIC value on the test data over all folds to generate the final network.

#### iRafNet

We use *iRafNet* to infer networks using prior information (it is not possible to run it without specifying priors). We called the *iRafNet* method of the package, setting the parameters *ntrees = 1000, mtry = round(sqrt(ncol(data)-1)),* and *npermut = 5* using the data filtered for missing values (see above) and then used the *Run_permutation* method with the same parameters. The final network was extracted using the *iRafNet_network* method by supplying the output of the previous method calls and setting the FDR cutoff parameter *TH = 0.05.* We used a custom implementation of *iRafNet* adjusted to make use of multiple CPUs which we made available at https://github.com/jhawe/irafnet_custom.

### Method evaluation via simulation study and cross cohort replication

To identify the inference method best suited for our application, we evaluated all described network inference methods independently on the simulated data as to 1) their ability to reconstruct the underlying ground truth network as well as 2) their robustness to noise in the supplied prior information. We further compared networks inferred independently on the different cohort data to assess stability of the network inference across different, yet similar, data. Performance was measured in terms of Matthew’s Correlation Coefficient (MCC) [29, 47, 106] between the inferred networks and the respective ground truth (simulation study) and the inferred networks on the different cohorts (cross cohort replication). It is defined as:

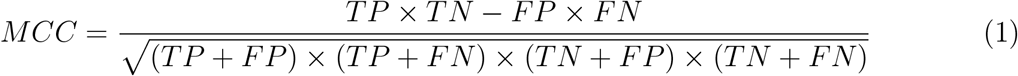

MCC was calculated using the *compare()* method as implemented in the *BDgraph* package (version 2.61).

### Transcription factor activities

We calculated transcription factor activities for all TFs extracted from the ReMap/ENCODE (see above) using the *plsgenomics* R package’s *T F A.estimate()* method (version 1.5-2) [107]. As input, we used the full expression matrix from KORA and LOLIPOP individually as well as the TFBS information encoded as an incidence matrix indicating for each TF its target genes. Target genes were defined as genes with an TFBS within their promoter region (2,000bp upstream and 1,000bp downstream of the TSS).

### Network prioritization and final network creation

Networks were inferred for each of the 107 meQTL and 444 eQTLGen *trans* hotspots with *gLASSO_P_* and *BDgraph_P_*, yielding networks with a median number of 67 and 20 edges for *gLASSO_P_* and 72 and 27 for *BDgraph_P_* over all hotspots, respectively. We filtered and ranked the networks based on the following criteria.

#### GWAS filtering

We filtered genetic loci with hits in genome-wide association studies (GWAS) using the current version (v1.0.2) of the GWAS catalog [52]. We extracted high LD (>0.8) SNPs and SNP aliases using the SNiPA tool [53] for each hotspot SNP. If any of the extracted SNP rsIDs had a match in the GWAS catalog, the hotspot’s inferred network was permitted for downstream analysis.

#### Network ranking

We utilized a self devised graph score for prioritizing final models for further investigation. The graph score reflects desirable biological properties, which can be assumed for the networks underlying the *trans* -QTL hotspots. The score is formulated such that 1) the adjacency of SNP-genes and SNPs is rated positively, 2) the presence of trans entities is rated positively if they are not connected directly to the SNP and 3) high graph density is rated negatively (i.e. sparser graphs yield higher scores). Specifically, the graph score *S_G_* for an inferred graph *G* is defined as:

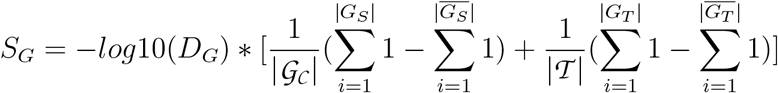

where: *D_G_* is the graph density, 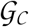 is the set of all SNP-Genes, 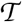 is the set of all *trans* entities, *G_S_* is the set of all SNP-genes adjacent to the SNP in *G* or directly connected to another SNP-Gene, 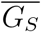 is the set of SNP-Genes in *G* but not connected directly to the SNP or one of the other SNP-Genes, *G_T_* is the set of trans entities in *G* which can be reached from any SNP-Gene without traversing the SNP or another *trans* gene first and 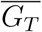 is the set of *trans* genes directly connected to the SNP. Only the cluster containing the SNP, i.e. the SNP itself and any nodes reachable from the SNP via any path in G, is considered for calculating *S_G_*; if the SNP is not present or no SNP gene has been selected in the final graph the score is set to 0.

In addition to the graph score, we ranked networks according to the total number of edges and nodes to prioritize smaller networks for detailed analysis.

#### Graph merging

Finally, we constructed hotspot networks containing only high confidence edges by merging the individually obtained networks from the two cohorts (KORA and LOLIPOP) and keeping only edges and nodes present in both networks. Nodes without any adjacent edges are not included in the final graph.

### Priors for skeletal muscle tissue

We downloaded Muscle tissue eQTL generated by Scott *et al.* (2016) [79] from https://theparkerlab.med.umich.edu/data/papers/doi/10.1038/ncomms11764/ and used local FDRs calculated from the provided p-values to define SNP-Gene priors. Gene expression data for Muscle tissue were obtained from the ARCHS^4^ [41] database. We downloaded all relevant Muscle expression data using the keywords “Skeletal_Muscle” with the ARCHS4 loader^17^ (N=194 samples). Expression data were normalized using the *ComBat* method implemented in the *sva* R package, providing dataset series ID as batch parameter.

#### TFBS prediction for muscle tissue

We used *factorNet* [80] to predict transcription factor binding sites from DNAse-seq chromatin accessibility data obtained from muscle cell lines. First, we trained a *factorNet* model for all TFs available for the K562 cell-line in ReMap [49]. ReMap ChIP-seq peaks functioned as a ground truth during training, DNAse-seq data from ENCODE^18^ [50, 51] and DNA sequence information formed the inputs. We downloaded DNAse-seq data for the LHCN-M2 muscle cell-line from ENCODE in bigWig format for hg38^19^. *FactorNet* was then run with default parameters, using as input 1) the DNA sequence and 2) the bigWig DNAse track for each of the trained ChIP-seq tanscription factors (N=179 TFs from ReMap). High confidence TFBS were extracted by setting a *factorNet* score cutoff of 0.999, merging overlapping regions and then retaining only regions with a *width* < *W*_0.95_, where *W*_0.95_ is the 95th percent quantile of the widths of all obtained regions.

### Colocalization analysis

GWAS summary statistics for schizophrenia were identified using the GWAS Atlas [108]^20^ and downloaded from http://walters.psycm.cf.ac.uk/clozuk_pgc2.meta.sumstats.txt.gz. Whole-blood *trans* -eQTL summary statistics for all SNP-Gene pairs from eQTLgen were downloaded from the eQTLgen website^21^. We used *fastENLOC* [76, 109]^22^ to calculate colocalization probabilities as described in the *fastENLOC* Github README using default parameters. To generate probabilistic eQTL annotations, we used *DAP-G* [110, 111]^23^ and created PIP files as needed using *TORUS* [112]^24^. For LD block definition, we utilized data available from LDetect [113]^25^.

### Software environment

In case no other information is given above, all calculations were performed using standard Unix commands and version 3.5.2 of the R statistical computing language^26^ on a Centos 7 Unix system. The Docker image used in this project is available from dockerhub at https://hub.docker.com/repository/docker/jhawe/r3.5.2_custom. The workflows for both the cohort and the simulation studies were implemented in Snakemake [114] and can be found on Github at https://github.com/jhawe/bggm. All calculations performed to arrive at the discussed results in this article can be obtained using the code in the pipeline. Data to run the workflow can be made available upon reasonable request by the authors.

## Declarations

### Availability of data and material

#### Data

All public data information and the respective sources are given in the methods section, including URLs for downloading the data where possible. The meQTL associations from Hawe et al. were directly obtained from the supplementary table 3 of the paper [10] and eQTLGen *trans* -eQTL directly from the eQTLGen browser^27^. The lists of derived hotspots for both data sets are made available in the supplement of this paper. Cohort data can be made available upon reasonable request by the authors.

#### Code

The complete code used in this project is provided via Github at https://github.com/jhawe/bggm. The analyses were implemented in the form of a Snakemake pipeline [114]. The software environment used to calculate the results is available as a Docker image via docker hub at https://hub.docker.com/repository/docker/jhawe/r3.5.2_custom, the corresponding dockerfile is available at the project’s Github repository.

### Ethics approval and consent to participate

All KORA participants have given written informed consent and the study was approved by the Ethics Committee of the Bavarian Medical Association. The LOLIPOP study is approved by the National Research Ethics Service (07/H0712/150) and all participants gave written informed consent.

### Consent for publication

KORA project agreement for this study was granted under K141/15g. The views expressed are those of the author(s) and not necessarily those of the Imperial College Healthcare NHS Trust, the NHS, the NIHR or the Department of Health.

### Authors’ contributions

MH conceived the study, JH performed the analyses. AB and AS assisted with use of GTEx v8 data. AB and FT contributed to the design of the data analysis strategy. CG, MW, KS, CH, SK, SW, HP, HG, AP, and MM provided KORA cohort data and JC the LOLIPOP data. JH and MH wrote the manuscript with input from all authors. All authors read and approved the final version of the manuscript.

## Acknowledgements

We thank the participants and research staff of LOLIPOP who made the study possible. The KORA-Study Group consists of A. Peters (speaker), J. Heinrich, R. Holle, R. Leidl, C. Meisinger, K. Strauch and their co-workers, who are responsible for the design and conduct of the KORA studies. We gratefully acknowledge the contribution of all members of field staff conducting the KORA study. Finally, we are grateful to all study participants of KORA for their invaluable contributions to this study.

## Footnotes

1 according to SNiPA: https://snipa.helmholtz-muenchen.de/snipa3/, [53]

2 https://ldlink.nci.nih.gov/?tab=ldmatrix

3 https://www.gtexportal.org/

4 https://www.encodeproject.org/

5 obtained from https://eqtlgen.org/trans-eqtls.html

6 file GTEx_Analysis_v8_trans_eGenes_fdr05.txt

7 http://tagc.univ-mrs.fr/remap/download/All/filPeaks_public.bed.gz

8 http://hgdownload.cse.ucsc.edu/goldenPath/hg19/encodeDCC/wgEncodeRegTfbsClustered/wgEncodeRegTfbsClusteredWithCellsV3.bed.gz

9 https://downloads.thebiogrid.org/Download/BioGRID/Release-Archive/BIOGRID-3.5.166/BIOGRID-ORGANISM-3.5.166.tab2.zip

10 https://storage.googleapis.com/gtex_analysis_v6p/rna_seq_data/GTEx_Analysis_v6p_RNA-seq_RNA-SeQCv1.1.8_gene_rpkm.gct.gz

11 file Whole_Blood_Analysis.v6p.all_snpgene_pairs.txt.gz from https://www.gtexportal.org/home/datasets

12 https://www.gtexportal.org/home/datasets

13 obtained from https://egg2.wustl.edu/roadmap/web_portal/chr_state_learning.html

14 https://cran.r-project.org/

15 https://www.bioconductor.org/

16 see also https://bioconductor.org/packages/release/bioc/vignettes/GENIE3/inst/doc/GENIE3.html

17 https://github.com/jhawe/archs4_loader

18 dataset ENCFF971AHO

19 dataset *ENCFF639MPM*

20 https://atlas.ctglab.nl/

21 https://www.eqtlgen.org/trans-eqtls.html, file ‘Full trans-eQTL summary statistics’

22 https://github.com/xqwen/fastenloc

23 https://github.com/xqwen/dap/

24 https://github.com/xqwen/torus

25 https://bitbucket.org/nygcresearch/ldetect-data/src/master/

26 https://www.r-project.org/

27 https://eqtlgen.org/trans-eqtls.html

