## Additional File 1 for "Network reconstruction for trans acting genetic loci using multi-omics data and prior information"

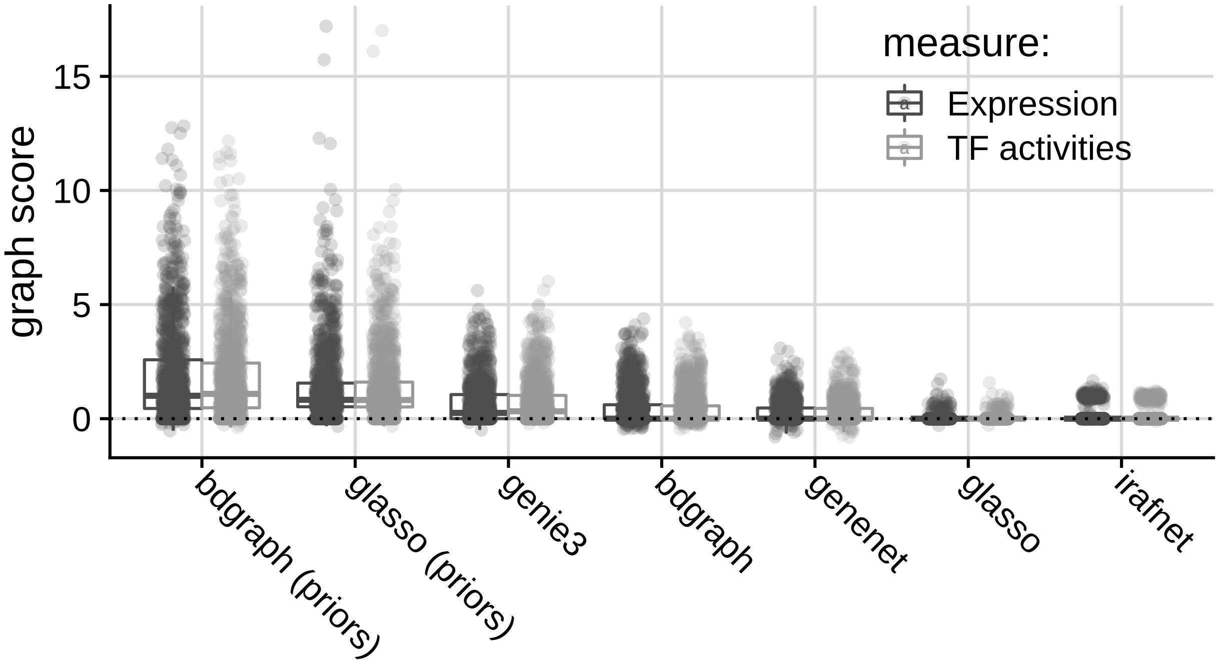


Supplementary Figure S 1: Overview over calculated graph scores over all hotspots for TFA based (light gray) and gene expression based (dark gray) network inference. y-axis shows the custom graph score, where higher scores indicate more desirable graph configurations. x-axis shows different inference methods. Boxplots show medians (horizontal line) and first and third quartiles (lower/upper box borders). Whiskers show $1.5 * IQR$ (inter-quartile range).

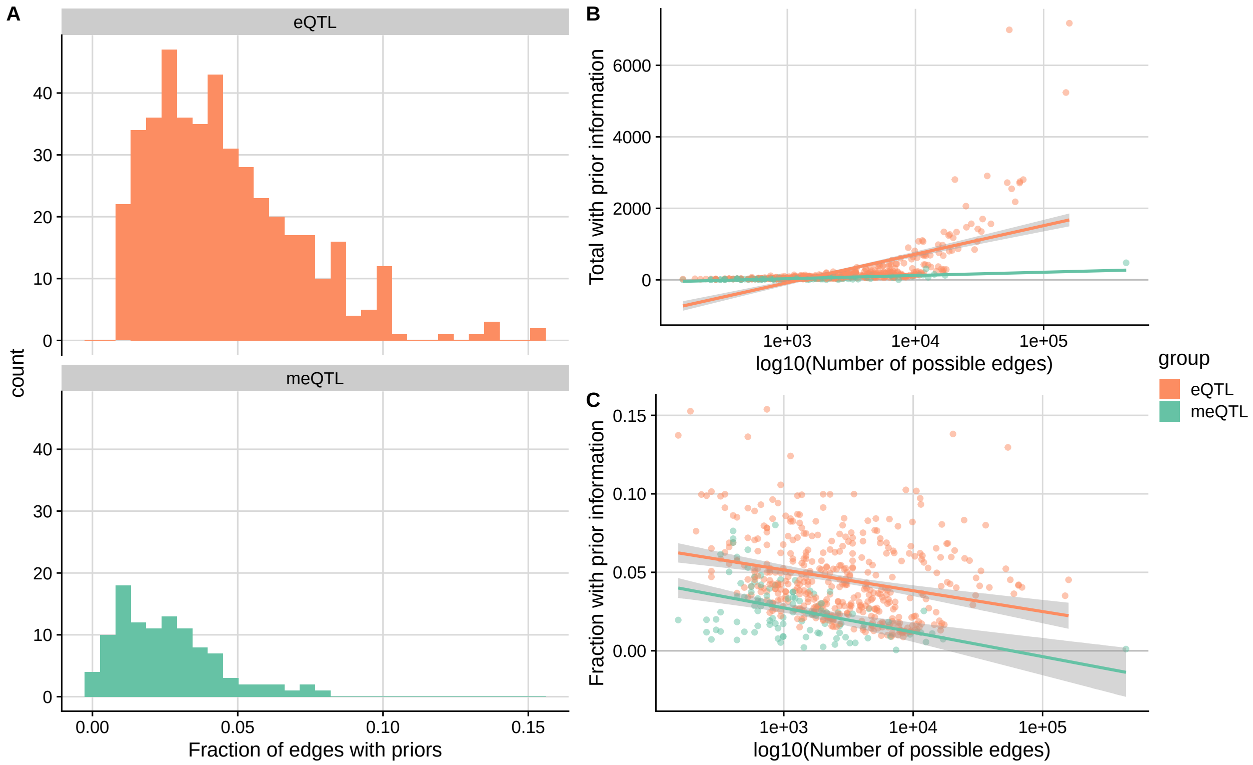


Supplementary Figure S 2: Overview over the number of collected priors for all *trans* hotspots. A) Histograms of the fraction of edges with annotated prior information. B) Total number of collected priors per hotspot (y-axis) plotted against the total number of possible edges (x-axis, log-scale). The total number of edges with prior information increases as the edge space increases. C) Same as B), but showing the fraction of edges with prior information on the y-axis. The fraction decreases as the total number of possible edges increases. Colors indicated QTL type (orange: eQTL, green: meQTL). Fitted lines represent the line of best fit with a linear model, grey areas show standard errors.

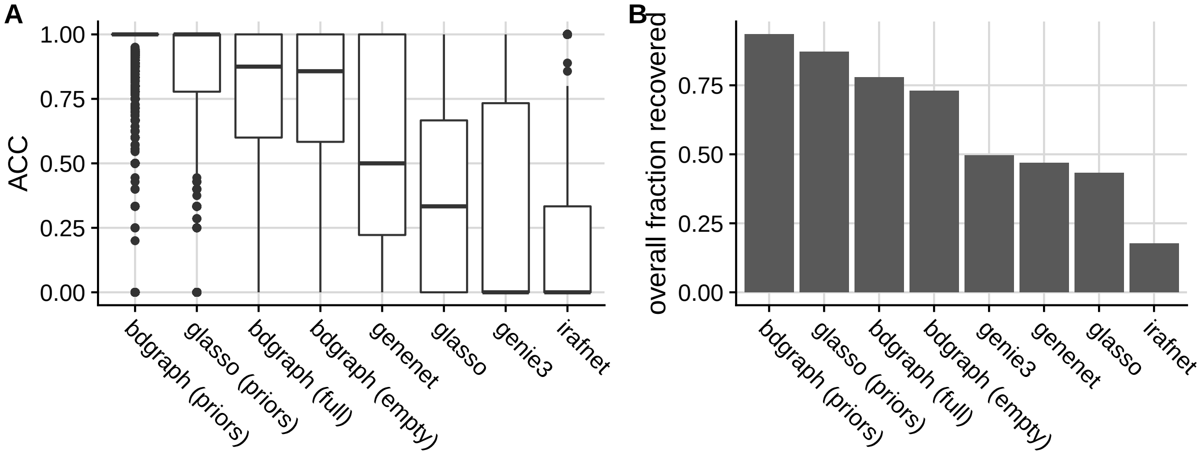


Supplementary Figure S 3: Performance of inference methods in recovering SNP links (edges between discrete and continuous data types) in the simulation. A) Boxplots showing fraction of recovered SNP associations (y-axis) for each method (x-axis) over all simulations. B) Summarized fractions, where for each methods the total number of recovered SNP associations over the total number of SNP associations in the generated ground truth graphs is calculated. Boxplots show medians (horizontal line) and first and third quartiles (lower/upper box borders). Whiskers show $1.5 * IQR$ (inter-quartile range), dots indicate outliers.

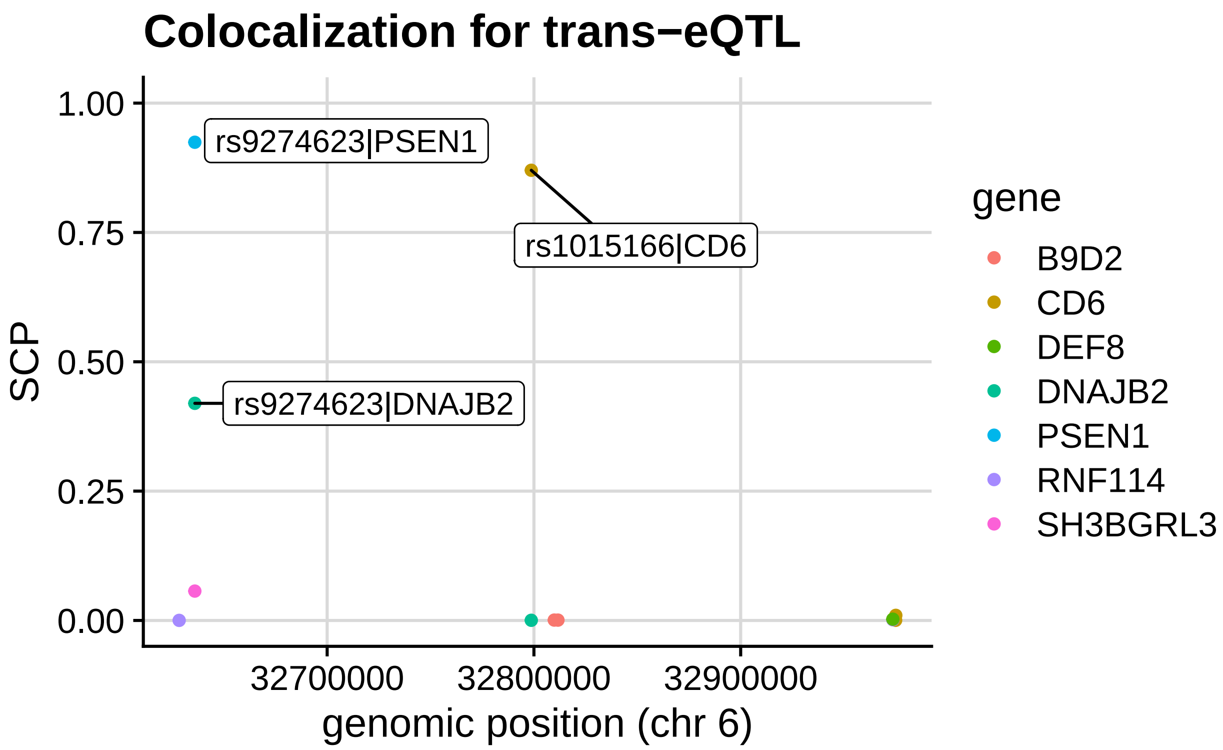


Supplementary Figure S 4: Results for the colocalization enrichment analysis for the schizophrenia locus. Y-axis shows the SNP-level colocalization probability for individual SNP-Gene pairs, x-axis genomic position of the respective SNPs.

| | sentinel | chr | cis_gene | gene_start | gene_end | gene_strand | gene_biotype |
|---|---|---|---|---|---|---|---|
| 1 | rs10870226 | chr10 | TTC40 | 134621896 | 134756327 | - | protein_coding |
| 2 | rs10870226 | chr10 | RP13-137A17.4 | 134757471 | 134778793 | - | lincRNA |
| 3 | rs10870226 | chr10 | RP13-137A17.5 | 134774844 | 134775741 | - | lincRNA |
| 4 | rs10870226 | chr10 | RP13-137A17.6 | 134779038 | 134789858 | + | lincRNA |
| 5 | rs17420384 | chr2 | AC105393.1 | 388412 | 416885 | + | lincRNA |
| 6 | rs17420384 | chr2 | AC105393.2 | 421057 | 422303 | + | lincRNA |
| 7 | rs17420384 | chr2 | AC093326.1 | 490944 | 492655 | - | lincRNA |
| 8 | rs17420384 | chr2 | AC093326.2 | 545805 | 546667 | + | lincRNA |
| 9 | rs17420384 | chr2 | AC093326.3 | 558204 | 578145 | + | lincRNA |
| 10 | rs2295981 | chr13 | LINC00354 | 112554299 | 112555490 | + | lincRNA |
| 11 | rs2295981 | chr13 | AL136302.1 | 112563079 | 112563148 | - | miRNA |
| 12 | rs2685252 | chr2 | AC105393.1 | 388412 | 416885 | + | lincRNA |
| 13 | rs2685252 | chr2 | AC105393.2 | 421057 | 422303 | + | lincRNA |
| 14 | rs2685252 | chr2 | AC093326.1 | 490944 | 492655 | - | lincRNA |
| 15 | rs2685252 | chr2 | AC093326.2 | 545805 | 546667 | + | lincRNA |
| 16 | rs2685252 | chr2 | AC093326.3 | 558204 | 578145 | + | lincRNA |
| 17 | rs57743634 | chr5 | SDHAP3 | 1568637 | 1594735 | - | pseudogene |
| 18 | rs57743634 | chr5 | CTD-2012J19.3 | 1594741 | 1611582 | + | lincRNA |
| 19 | rs57743634 | chr5 | CTD-2012J19.2 | 1598242 | 1598362 | + | pseudogene |
| 20 | rs57743634 | chr5 | RP11-43F13.1 | 1599035 | 1634120 | - | pseudogene |
| 21 | rs57743634 | chr5 | CTD-2012J19.1 | 1614951 | 1616449 | + | pseudogene |
| 22 | rs57743634 | chr5 | MIR4277 | 1708900 | 1708983 | - | miRNA |
| 23 | rs57743634 | chr5 | CTD-2587M23.1 | 1725264 | 1728287 | + | lincRNA |

Supplementary Table S 1: Sentinels and their annotated cis genes removed from analysis due to the genes not being measured on the microarrays. Sentinels rs1570038 and rs7924137 did not have any cis genes annotated.

| method | R=0 | R=0.1 | R=0.2 | R=0.3 | R=0.4 | R=0.5 | R=0.6 | R=0.7 | R=0.8 | R=0.9 | R=1 | R=rbinom |
|---|---|---|---|---|---|---|---|---|---|---|---|---|
| **bdgraph (priors)** | **0.93** | **0.91** | **0.87** | 0.83 | 0.80 | 0.77 | 0.72 | 0.69 | 0.64 | 0.60 | 0.55 | **0.83** |
| **glasso (priors)** | 0.87 | 0.81 | 0.74 | 0.66 | 0.60 | 0.53 | 0.46 | 0.41 | 0.34 | 0.27 | 0.21 | 0.42 |
| **bdgraph (empty)** | 0.84 | 0.84 | 0.84 | **0.84** | **0.84** | **0.84** | **0.84** | **0.85** | **0.85** | **0.85** | **0.85** | **0.83** |
| **bdgraph (full)** | 0.84 | 0.84 | 0.84 | **0.84** | **0.84** | **0.84** | **0.84** | **0.85** | **0.84** | **0.84** | **0.85** | **0.83** |
| **genenet** | 0.65 | 0.66 | 0.65 | 0.65 | 0.65 | 0.66 | 0.66 | 0.67 | 0.67 | 0.67 | 0.67 | 0.65 |
| **irafnet** | 0.45 | 0.43 | 0.42 | 0.39 | 0.37 | 0.36 | 0.34 | 0.31 | 0.28 | 0.24 | 0.20 | 0.42 |
| **glasso** | 0.43 | 0.43 | 0.43 | 0.43 | 0.43 | 0.43 | 0.44 | 0.44 | 0.44 | 0.45 | 0.46 | 0.42 |
| **genie3** | 0.38 | 0.37 | 0.37 | 0.37 | 0.37 | 0.38 | 0.38 | 0.39 | 0.39 | 0.38 | 0.40 | 0.35 |

Supplementary Table S 2: Table showing an overview over the performance (mean MCC) in the simulation study for each method for all prior noise scenarios, sorted by first column. Highest mean MCC for each scenario is indicated in bold.

| method | R=0 | R=0.1 | R=0.2 | R=0.3 | R=0.4 | R=0.5 | R=0.6 | R=0.7 | R=0.8 | R=0.9 | R=1 | R=rbinom |
|---|---|---|---|---|---|---|---|---|---|---|---|---|
| **bdgraph (priors)** | **0.93** | **0.91** | **0.87** | **0.84** | 0.81 | 0.77 | 0.73 | 0.70 | 0.65 | 0.61 | 0.56 | 0.82 |
| **glasso (priors)** | 0.87 | 0.81 | 0.74 | 0.66 | 0.60 | 0.54 | 0.47 | 0.42 | 0.35 | 0.29 | 0.22 | 0.42 |
| **bdgraph (empty)** | 0.84 | 0.84 | 0.84 | **0.84** | **0.84** | **0.84** | **0.84** | **0.84** | **0.84** | **0.84** | **0.84** | **0.83** |
| **bdgraph (full)** | 0.84 | 0.84 | 0.84 | **0.84** | **0.84** | **0.84** | **0.84** | **0.84** | **0.84** | **0.84** | **0.84** | **0.83** |
| **genenet** | 0.63 | 0.64 | 0.63 | 0.63 | 0.63 | 0.64 | 0.64 | 0.65 | 0.65 | 0.65 | 0.65 | 0.63 |
| **glasso** | 0.43 | 0.43 | 0.43 | 0.43 | 0.43 | 0.43 | 0.43 | 0.44 | 0.44 | 0.44 | 0.45 | 0.42 |
| **irafnet** | 0.37 | 0.35 | 0.34 | 0.31 | 0.29 | 0.28 | 0.26 | 0.23 | 0.21 | 0.18 | 0.16 | 0.40 |
| **genie3** | 0.31 | 0.30 | 0.30 | 0.30 | 0.30 | 0.31 | 0.31 | 0.32 | 0.32 | 0.31 | 0.31 | 0.28 |

Supplementary Table S 3: Same as ST 2, but showing mean F1 scores instead of MCC. Highest mean F1 for each scenario is indicated in bold.

| method | N= 50 | 100 | 150 | 200 | 250 | 300 | 350 | 400 | 450 | 500 | 550 | 600 |
|---|---|---|---|---|---|---|---|---|---|---|---|---|
| **bdgraph (priors)** | **0.86** | **0.86** | **0.87** | **0.89** | **0.90** | **0.91** | **0.92** | **0.92** | **0.92** | **0.93** | **0.93** | **0.93** |
| **glasso (priors)** | 0.83 | 0.85 | 0.86 | 0.86 | 0.87 | 0.87 | 0.87 | 0.87 | 0.88 | 0.88 | 0.88 | 0.88 |
| **bdgraph (empty)** | 0.43 | 0.57 | 0.64 | 0.69 | 0.74 | 0.76 | 0.78 | 0.80 | 0.81 | 0.82 | 0.83 | 0.84 |
| **bdgraph (full)** | 0.42 | 0.56 | 0.64 | 0.69 | 0.74 | 0.76 | 0.78 | 0.80 | 0.81 | 0.82 | 0.83 | 0.84 |
| **irafnet** | 0.32 | 0.38 | 0.41 | 0.42 | 0.43 | 0.44 | 0.44 | 0.44 | 0.45 | 0.45 | 0.45 | 0.45 |
| **genenet** | 0.29 | 0.43 | 0.50 | 0.54 | 0.57 | 0.60 | 0.61 | 0.63 | 0.63 | 0.65 | 0.65 | 0.66 |
| **genie3** | 0.26 | 0.30 | 0.32 | 0.33 | 0.34 | 0.34 | 0.35 | 0.35 | 0.36 | 0.36 | 0.36 | 0.36 |
| **glasso** | 0.20 | 0.27 | 0.31 | 0.34 | 0.37 | 0.38 | 0.39 | 0.40 | 0.41 | 0.42 | 0.42 | 0.43 |

Supplementary Table S 4: Table showing an overview over the performance (mean MCC) in the simulation study for each method for different sub samplings of simulated data (increasing from left to right), sorted by first column. Highest mean MCC for each scenario is indicated in bold.



| method | N= 50 | 100 | 150 | 200 | 250 | 300 | 350 | 400 | 450 | 500 | 550 | 600 |
|---|---|---|---|---|---|---|---|---|---|---|---|---|
| **bdgraph (priors)** | **0.86** | **0.85** | **0.87** | **0.89** | **0.90** | **0.91** | **0.91** | **0.92** | **0.92** | **0.93** | **0.93** | **0.93** |
| **glasso (priors)** | 0.83 | **0.85** | 0.86 | 0.86 | 0.87 | 0.87 | 0.87 | 0.87 | 0.88 | 0.88 | 0.88 | 0.88 |
| **bdgraph (empty)** | 0.42 | 0.55 | 0.63 | 0.68 | 0.73 | 0.75 | 0.78 | 0.79 | 0.81 | 0.82 | 0.83 | 0.84 |
| **bdgraph (full)** | 0.41 | 0.55 | 0.63 | 0.68 | 0.73 | 0.75 | 0.78 | 0.79 | 0.81 | 0.82 | 0.83 | 0.84 |
| **genenet** | 0.24 | 0.39 | 0.46 | 0.51 | 0.55 | 0.57 | 0.59 | 0.60 | 0.61 | 0.63 | 0.63 | 0.64 |
| **irafnet** | 0.24 | 0.30 | 0.32 | 0.34 | 0.34 | 0.35 | 0.36 | 0.36 | 0.36 | 0.37 | 0.37 | 0.37 |
| **glasso** | 0.19 | 0.27 | 0.31 | 0.34 | 0.36 | 0.38 | 0.39 | 0.40 | 0.41 | 0.42 | 0.42 | 0.43 |
| **genie3** | 0.19 | 0.23 | 0.24 | 0.26 | 0.26 | 0.27 | 0.28 | 0.28 | 0.28 | 0.29 | 0.29 | 0.29 |

Supplementary Table S 5: Table showing an overview over the performance (mean F1) in the simulation study for each method for different sub samplings of simulated data (increasing from left to right), sorted by first column. Highest mean F1 for each scenario is indicated in bold.

| | method | Expression | TF activities |
|---|---|---|---|
| 1 | glasso (priors) | 0.74 (0.18) | **0.75 (0.18)** |
| 2 | bdgraph (priors) | 0.46 (0.1) | **0.49 (0.1)** |
| 3 | irafnet | 0.36 (0.23) | **0.43 (0.21)** |
| 4 | genenet | 0.31 (0.12) | **0.33 (0.12)** |
| 5 | bdgraph (empty) | 0.29 (0.11) | **0.3 (0.12)** |
| 6 | glasso | 0.25 (0.21) | **0.28 (0.22)** |
| 7 | genie3 | **0.2 (0.23)** | **0.2 (0.19)** |

Supplementary Table S 6: Table shows the mean cross cohort replication MCC for expression and TF activity based analyses for each method and standard deviations in parentheses. Highest mean MCC per method is indicated in bold.
